# Dissecting Mammalian Spermatogenesis Using Spatial Transcriptomics

**DOI:** 10.1101/2020.10.17.343335

**Authors:** Haiqi Chen, Evan Murray, Anisha Laumas, Jilong Li, Xichen Nie, Jim Hotaling, Jingtao Guo, Bradley R. Cairns, Evan Z. Macosko, C. Yan Cheng, Fei Chen

## Abstract

Single-cell RNA sequencing has revealed extensive molecular diversity in gene programs governing mammalian spermatogenesis but fails to delineate their dynamics in the native context of seminiferous tubules — the spatially-confined functional units of spermatogenesis. Here, we use Slide-seq, a novel spatial transcriptomics technology, to generate a comprehensive spatial atlas that captures the spatial gene expression patterns at near single-cell resolution in the mouse and human testis. By using Slide-seq data, we devise a computational framework that accurately localizes testicular cell types in individual seminiferous tubules. Unbiased spatial transcriptome analysis systematically identifies spatially patterned genes and gene programs, nominating genes with previously underappreciated but important functions in spermatogenesis. Using the human testicular spatial atlas, we identify two spatially segregated spermatogonial populations composed of stem cells at distinct transcriptional states. Finally, a comparison of the spatial atlas generated from the wild type and diabetic mouse testis reveals a disruption in the spatial cellular organization in diabetic seminiferous tubules.

## Introduction

Spermatogenesis — the biological process of sperm production — plays a crucial role in controlling male fertility. However, our understanding on the molecular basis underlying spermatogenesis remains largely incomplete. This is due to a lack of experimental approaches that can faithfully recapitulate spermatogenesis *in vitro* in spite of recent advances in three-dimensional germ cell cultures (Mahmoud, 2012; Alves-Lopes and Stukenborg, 2018), as well as a lack of tools to systematically profile spermatogenesis *in vivo*.

High-throughput molecular profiling technologies such as single-cell RNA sequencing (scRNA-seq) provides an attractive alternative to study spermatogenesis as it captures the heterogeneity in gene expression profiles of germ cells at each stage of development (Green et al., 2018; Lukassen et al., 2018; Wang et al., 2018; Guo et al., 2018; Hermann et al., 2018). However, scRNA-seq fails to profile developing germ cells in the native context of a seminiferous tubule — the spatially-confined functional unit of spermatogenesis — due to cell dissociation. The difficulty of studying spermatogenesis using scRNA-seq is further compounded by somatic cell types co-existing with the germ cells in the testis (Chen et al., 2017; Griswold, 2018; Smith and Walker, 2014). Failure of scRNA-seq to capture the spatial interaction between the germ cell lineage and the somatic cell lineage impedes a comprehensive understanding of spermatogenesis. Previous methods for extracting spatial transcriptomic information from the tissues such as individual-cell laser-capture microdissection or multiplexed in situ hybridization are low throughput and require prior knowledge of the cell types or the genes to be targeted (Jan et al., 2017; Shalek and Satija, 2015). Therefore, an unbiased, high-throughput molecular profiling method capable of capturing the spatial context of testicular cells at high resolution is needed to truly recapitulate spermatogenesis *in vivo*.

We have recently developed Slide-seq, a spatial transcriptomics technology that enables high-throughput spatial genomics at 10-μm resolution (Rodriques et al., 2019; Stickels et al., 2020). We applied Slide-seqV2 to both adult mouse and adult human testis samples and developed an algorithmic pipeline to build a spatial transcriptome atlas for mammalian spermatogenesis. This atlas spatially localized testicular cell types within seminiferous tubules with high accuracy. Using this spatial atlas, we systematically identified spatially patterned (SP) genes with distinct biological functions for both the mouse and the human testis. Among those SP genes, we discovered *Habp4* (hyaluronan binding protein 4) as a potential novel regulator of chromatin remodeling in developing germ cells. Furthermore, we used the mouse spatial atlas to capture the stage-dependent gene expression patterns of somatic cell type Leydig cells and macrophages. Moreover, in human testes, we provided evidence that two spatially distinct spermatogonial stem cell (SSC) neighborhoods co-exist in a seminiferous tubule. Finally, we demonstrated a disruption in the spatial structure of seminiferous tubules as a prevalent phenotype in diabetic testes using a leptin-deficient model. Together, we have demonstrated the power of high-resolution spatial transcriptomics in revealing novel molecular information of mammalian spermatogenesis. This rich source of high resolution spatial transcriptomic data and its associated computational data analysis pipelines will be provided for the community to further efforts in developing analytical approaches for *in situ* molecular profiling, spatial mapping of cell types and tissue structures, as well as characterizing the functional dynamics of gene programs.

## Results

### Building a Spatial Transcriptome Atlas for Mouse Spermatogenesis

A workflow was established to enable the capture of testicular mRNAs onto Slide-seq arrays (Fig 1A & 1B). Each array is composed of 10-μm beads with unique spatial barcodes which are sequenced *in situ* to assign each bead to a unique spatial location (Fig 1A). Next, a thin frozen testis cross-section (~10 μm) was placed on top of the spatially indexed bead array during cryosectioning, and mRNA from the testis section was captured by the beads. Sequencing of subsequent barcoded libraries can be uniquely matched to the spatial coordinates on the bead array (Fig 1B). The Slide-seq array diameter (3 mm) allowed for a full coverage of an adult mouse testis cross-section. Using the established workflow, we routinely obtained a high mRNA capture rate at an average of 784 ± 25 (mean ± SEM, n= 3 experiments) transcripts (Unique Molecular Identifiers, or UMIs) per bead with an average total number of beads at 29333 ± 1514 (mean ± SEM, n= 3 arrays) per array. Mapping of testicular cell types onto Slide-seq beads using a non-negative matrix factorization regression (NMFreg)-based statistical method (Rodriques et al., 2019) and a scRNA-seq reference (Green et al., 2018) yielded cell-type assignments reflecting structures of seminiferous tubules (Fig 1C) and known positions of testicular cell types (Fig 1D). For example, beads assigned as elongating/elongated spermatids (ESs) were accurately assigned to the center of the seminiferous tubules whereas beads assigned as Leydig cells distributed as clusters in the interstitial space in between seminiferous tubules (Fig 1D). The accuracy of the cell type assignment was further confirmed by the enrichment of cell type marker genes in the corresponding cell type cluster (Fig S1).

**Figure 1.**
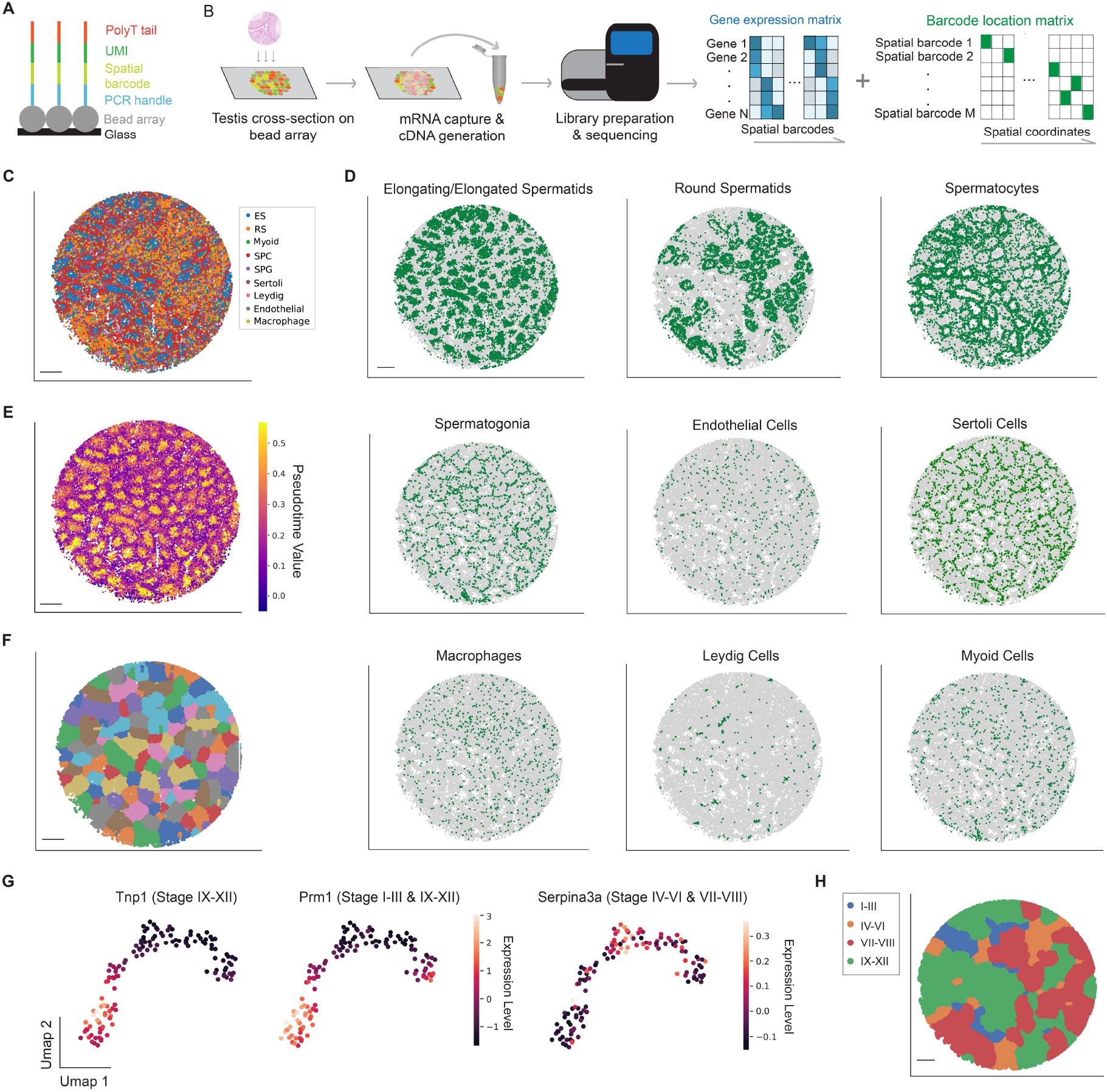
Establishment of the Mouse Testicular Spatial Transcriptome Atlas. (A) Sequence schematic of the Slide-seq bead oligonucleotides. (B) Slide-seq workflow for testicular samples. (C) Spatial mapping of testicular cell types. ES, elongating/elongated spermatid; RS, round spermatid; SPC, spermatocyte; SPG, spermatogonium. Scale bar, 300 μm. (D) Spatial mapping of individual testicular cell type. Scale bar, 300 μm. (E) Pseudotime reconstruction of germ cell developmental trajectory. Scale bar, 300 μm. (F) Digital segmentation of the seminiferous tubules. Scale bar, 300 μm. (G) Umap projection of the seminiferous tubules. Tubule clusters were colored by genes with known stage-specific expression patterns. (H) Spatial mapping of the four stage clusters. Scale bar, 300 μm.

Following the cell type assignment, we sought to assign information of the seminiferous tube and the stage of the seminiferous epithelium cycle to each bead. To this end, we first performed pseudotime analysis to rank each bead along a transcriptional trajectory (Methods). This analysis recapitulated the known germ cell developmental trajectory which originates from the basement membrane and migrates towards the lumen of the seminiferous tubules (Fig 1E). Next, we developed a computational pipeline to automatically group Slide-seq beads belonging to the same seminiferous tubule (Fig S2B, Methods) based on the observation that the reconstructed pseudotime image retained the morphological details of seminiferous tubule cross-sections (Fig S2A). Finally, a spermatogenic cycle can be divided into substages, with each stage containing a distinct association of germ cell subtypes (Clermont and Trott, 1969). By projecting the expression profiles of genes with known stage-dependent expression pattern (e.g., *Prm1*) onto the UMAP (uniform manifold approximation and projection) embedding of the segmented seminiferous tubules (Fig 1G), we assigned each tubule with one of the four major stage clusters (Stage I-III, IV-VI, VII-VIII, and IX-XII) (Fig 1H). Additional genes with known stage-specific expression patterns (Klaus et al., 2016) were used to validate the stage assignment. Together, we generated a spatial atlas for mouse spermatogenesis from the ground up by assigning each Slide-seq bead to its corresponding cell type, seminiferous tubule and stage.

### A Systematic Identification of Spatially Patterned (SP) Genes in Seminiferous Tubules

Testicular cell types are organized in a spatially segregated fashion at the level of seminiferous tubules. Therefore, genes with spatially non-random distribution may exert functions in different cell types or sub-cell types. To this end, we used the spatial transcriptome atlas to identify genes with spatially nonrandom distribution (hereafter referred as spatially patterned genes, or SP genes) at the level of individual seminiferous tubules. By applying a generalized linear spatial model-based statistical method (Sun et al., 2020) to 103 seminiferous tubules gathered from three normal mouse samples, we systematically identified 277, 702, 298, and 665 SP genes for stage I-III, IV-VI, VII-VIII, and IX-XII tubules, respectively (Table S1), with the majority of them being novel genes whose spatial expression patterns had not been previously characterized(Fig 2A). We confirmed the accuracy of the SP gene identification using single molecule fluorescence *in situ* hybridization (smFISH) on several novel genes: *Smcp* (sperm mitochondrial-associated cysteine-rich protein), shown by our analysis as a SP gene with high expression near the center of the seminiferous tubule, was found mainly expressed in elongating spermatids near the tubule lumen (Fig 2B); whereas the SP gene *Lyar* which encodes the cell growth-regulating nucleolar protein involved in processing of preribosomal RNA (Miyazawa et al., 2014) was enriched in spermatocytes near the basement membrane, consistent with our observation (Fig 2B). Moreover, the spatial expression patterns of some SP genes are restricted to only one of the four stage clusters (Table S1). For example, gene *Tnp1* (transition protein 1) were only highly expressed near the tubule lumen in a stage IX-XII tubule (Fig 2C), consistent with the previous finding that the expression of the TNP1 protein was restricted to mouse spermatids of steps 9–12 (Klaus et al., 2016).

**Figure 2.**
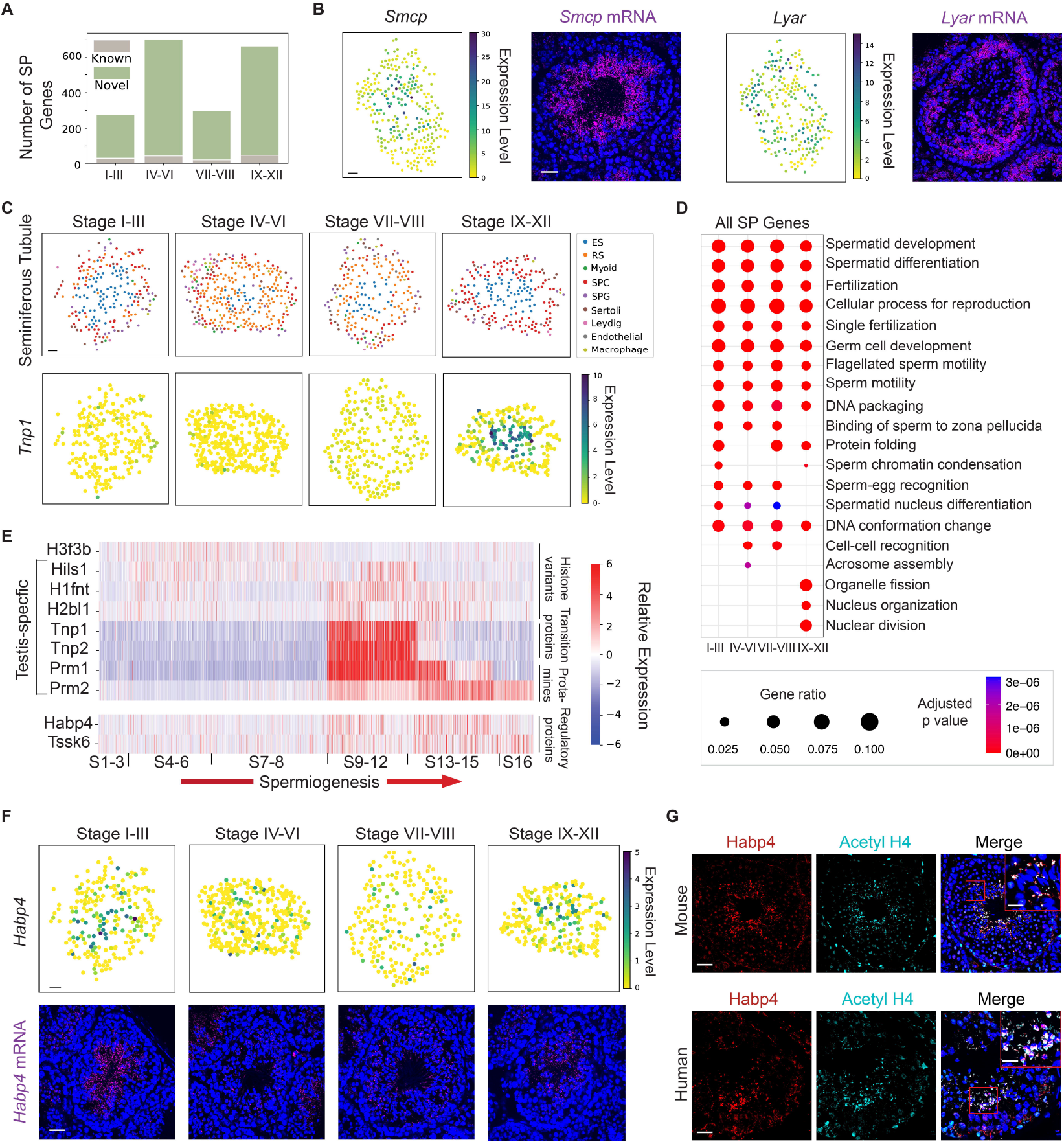
Systematic identification of spatially patterned (SP) genes in the seminiferous tubules. (A) The number of genes with previously known spatial pattern vs. the number of newly identified genes using the spatial transcriptome atlas in each stage of the seminiferous epithelium cycle. (B) The spatial expression pattern of *Smcp* and *Lyar* revealed by both the spatial transcriptome atlas and single-molecule fluorescence *in situ* hybridization (smFISH). Scale bar, 30 μm for the digitally reconstructed seminiferous tubule images and 50 μm for the smFISH images. (C) The spatial transcriptome atlas reveals the stage-dependent spatial expression pattern of *Tnp1*. ES, elongating/elongated spermatid; RS, round spermatid; SPC, spermatocyte; SPG, spermatogonium. (D) Gene ontology enrichment analysis on SP genes from the four stage clusters. (E) The temporal dynamics of SP genes enriched in nucleus organization during spermiogenesis. (F) The spatial transcriptome atlas and smFISH reveal the stage-dependent spatial expression pattern of *Habp4*. (G) Co-localization of Habp4 protein with the acetylated histone 4 (Acetyl H4) in mouse and human spermatids. Scale bar for the mouse images, 40μm; 10 μm for the inset. Scale bar for the human images, 40μm; 15 μm for the inset.

Next, we performed gene ontology (GO) enrichment analysis on the identified SP genes. Although signaling pathways such as spermatid development and differentiation were shared across all stages, we found that certain signaling pathways were stage-enriched (Fig 2D). For example, genes involved in acrosome assembly were enriched in stage IV-VI. And processes related to cell division (e.g., meiotic nuclear division) including genes such as the Sycp family were enriched in stage IX-XII (Fig 2E; Table S1), consistent with the observation that meiosis mainly takes place in stage IX-XII tubules.

In addition, the GO enrichment analysis identified a group of SP genes enriched in signaling pathways related to chromatin remodeling (e.g., DNA conformation change, DNA packaging and nucleus organization). During spermiogenesis—the final stage of spermatogenesis, post-meiosis spermatids undergo a remarkable chromatin remodeling process in which the majority of histones in haploid spermatids are replaced with protamines (O’Donnell, 2014; Bao and Bedford, 2016; Rathke et al., 2014). However, our understanding of genes participating in this process remains incomplete. To this end, we focused on SP genes enriched in the signaling pathways of DNA confirmation change, DNA packaging and nucleus organization. These genes included somatic histone component *H3f3b*; testis-specific histone variants *Hils1, H2bl1* (also known as *1700024P04Rik), H1fnt*; transition protein genes *Tnp1, Tnp2*; and protamine genes *Prm1, Prm2*. First using smFISH, we confirmed the predicted spatial localizations of a subset of these genes as their spatial expression patterns were previously unknown (Fig S3). Second, we looked at their gene expression profiles along the progression of spermiogenesis by extracting Slide-seq beads assigned as spermatids from the atlas and ordered them based on the stage they belonged to (Fig 2E). As expected, somatic histone variant *H3f3b* was highly expressed in early round spermatids (RSs), but its expression level decreased as spermiogenesis proceeded. A previous study suggested that the perturbation of *H3f3b* caused male infertility phenotypes (Tang et al., 2015). In contrast, testis-specific histone variants showed highest expression in elongating spermatids. *Hils1* expression decreased towards the late stages, similarly to the transition protein genes *Tnp1* and *Tnp2*. Of interest, the expression of *Tnp1* and *Tnp2* were restricted in a narrow temporal window along the developmental trajectory of spermatids, consistent with their transitional role of histone eviction and protamine recruitment (Meistrich et al., 2003; Shirley et al., 2004; Zhao et al., 2004). In contrast, the expression of *H1fnt* and *H2bl1* persisted towards the late stages of spermatid differentiation similar to *Prm1*, suggesting a role for these histone variants during the final stage of chromatin condensation. Finally, we captured a difference in timing for *Prm1* and *Prm2* to reach their peak expression level (Fig 2E), indicating a potential functional divergence of the two protamine genes. Taken together, we have successfully used spatial transcriptome data to recapitulate the temporal dynamics of histone-to-protamine transition during spermiogenesis.

### *Habp4* is Associated with Chromatin Remodeling in Spermatids

Two additional SP genes *Tssk6* and *Habp4* were also enriched in the signaling pathways related to chromatin remodeling. The expression of *Tssk6* and *Habp4* were both enriched in mouse elongating spermatids (Fig 2E). TSSK6, member of testis-specific serine/threonine kinases has been shown to be necessary for histone-to-protamine transition during spermiogenesis (Jha et al., 2017). However, the functional role of *Habp4* in the testis is unclear. To start unraveling the biological function of *Habp4*, we first profiled its spatial expression pattern across all stage clusters using the spatial atlas and smFISH (Fig 2F). We found that *Habp4* was highly expressed near the tubule lumen among the spermatid population, but only at stage I-III and IX-XII tubules (Fig 2F).

Next, we hypothesized that the similar temporal expression pattern of *Tssk6* and *Habp4* as shown in Fig 2E might also suggest functional similarities. Displacement of histones by transition proteins and protamines is accompanied by several histone post-translational modifications (PTMs). Biochemical studies indicated that histone acetylation, especially histone H4 acetylation (acetyl H4), facilitated the displacement of histones (Meistrich et al., 1992)(Awe and Renkawitz-Pohl, 2010). Immunofluorescence analysis showed co-localization of acetyl H4 with HABP4 proteins in elongating spermatids both in the mouse and human testis (Fig 2G), indicating a role of HABP4 in the chromatin remodeling process during histone-protamine transition. Moreover, DNA strand breaks (DSBs) have been observed in spermatocytes and spermatids and are proposed as a mechanism to facilitate DNA conformational changes during spermiogenesis (Marcon and Boissonneault, 2004). The active, phosphorylated form of H2AX (H2A histone family, member X; H2AFX), γH2AX, is present with the DSBs (Leduc et al., 2008). Although we observed minimal colocalization of HABP4 protein and γH2AX in the mouse testis, we found that these two proteins co-localized within a subset of spermatocytes in the human testis (Fig S4). This finding suggests a possible separate function of HABP4 in the human testis, likely associated with the formation or repair of DSBs.

### Temporal Expression Dynamics of X Chromosome-linked Escape Genes in Spermatids

Inspired by the approach of using the spatial atlas to infer the temporal dynamics of genes invovled in chromatin remodeling as described above, we sought to use the same approach to profile genes excpaing the meiotic sex chromosome inactivation (MSCI) (Daish and Grützner, 2010; Turner, 2007). A subgroup of sex chromosome-linked genes is able to escape MSCI in that these genes show increased expression after meiosis. Ranking the Slide-seq beads along the temporal axis of spermatogenesis allowed for the sensitive detection of spermatid-specific X chromosome-linked escape genes (Fig S5). These escape genes included previously annotated escape genes such as *Akap4, Prdx4, Pgrmc1, Tspan6, Eif1ax, Ctag2*, and *Cypt3*. Their reactivation has been shown to be dependent on a RNF8- and SCML2-mediated mechanism (Adams et al., 2018). Moreover, our data demonstrated that X-linked escape genes exhibited three distinct temporal expression patterns during spermatid development (Fig S5). The first pattern showed a peak expression in round spermatids directly following meiosis. The second pattern represented an upregulation of expression in escape genes only at the early stages of elongating spermatid development whereas the genes with the third expression pattern were upregulated throughout the entire developmental process of elongating spermatids. The different timing in the reactivation of these X-linked escape genes suggests their potential functional divergence in regulating spermatid development.

### Identifying the Stage-specific Expression Patterns of Leydig Cells and Macrophages

Following a detailed profiling of the genes with non-random spatial distributions using the spatial transcriptome atlas, we next turned to genes with stage-specific expression patterns in different testicular cell types. Stage-specific gene expression has been shown as a fundamental characteristic of germ cells as well as somatic cells such as Sertoli cells (Green et al., 2018; Johnston et al., 2008; Linder et al., 1991; Sugimoto et al., 2012). However, the stage-specific expression patterns of somatic cells in the interstitial and peritubular space remains largely unexplored. To this end, we used the spatial transcriptome atlas to perform differential gene expression analysis among Leydig cells and macrophages assigned to different stage clusters. We first identified stage-specific genes of Leydig cells in the clusters of I-III, IV-VI, and VII-VIII, respectively (Fig 3A). Four of these genes (*Ankar*, *Tekt2*, *1700011A15Rik*, and *Pdilt*) have been previously reported to be stage-dependently expressed in Leydig cells (Jauregui et al., 2018). To further validate the result, we used smFISH to confirm the stage-specificity of *1700017N19Rik* in stage IV-VI-associated Leydig cells (Fig 3B). Of interest, we noticed that Leydig cells expressing *1700017N19Rik* were preferentially localized close to the peritubular space (Fig 3B) and were in close spatial proximity with spermatogonia (Fig S6A, white arrow). Quantitative analysis of the spatial atlas data showed that the Euclidean distance from *1700017N19Rik*-expressing Leydig cell beads to the nearest spermatogonium beads was significantly shorter than that from Leydig cell beads with no *1700017N19Rik* expression to the neighboring spermatogonium beads (two-sided *t* test, p <0.05) (Fig S6A, bar plot). The sparsity and the spatial localization of the *1700017N19Rik*-expressing Leydig cells resembled those of the stem Leydig cells (Li et al., 2016). Indeed, we found that the *1700017N19Rik*-expressing Leydig cells also expressed stem Leydig cell marker *Nr2f2* (Fig 3C), indicating that *1700017N19Rik* may be involved in the regulation of stem leydig cell functions.

**Figure 3.**
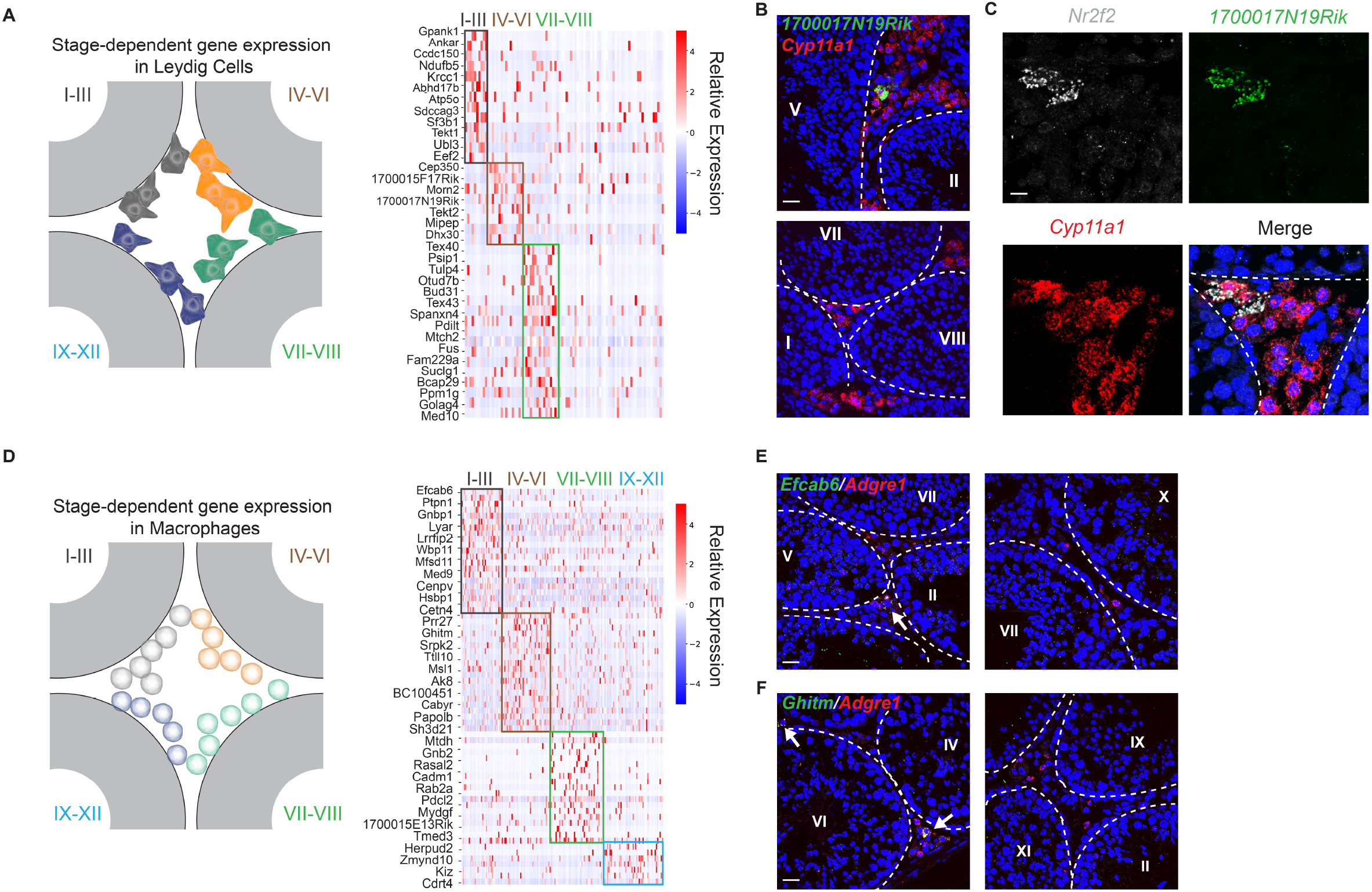
Stage-dependent gene expression patterns in Leydig cells and macrophages. (A) Left: schematic of the spatial localizations of Leydig cells. Right: genes exhibiting stage-dependent expression patterns in Leydig cells. (B) *17700017N19Rik*-expressing Leydig cells (marked by *Cyp11a1*) localize near a stage IV-VI seminiferous tubule, but not close to tubules at other stages. The basement membrane of the tubules are outlined by white dashed lines. Scale bar, 20 μm (C) smFISH experiments show that *1700017NI9Rik*-expressing Leydig cells also express stem Leydig cell marker *Nr2f2*. The basement membrane of the seminiferous tubules are outlined by white dashed lines. Scale bar, 10 μm (D) Left: schematic of the spatial localizations of macrophages. Right: genes exhibiting stage-dependent expression patterns in macrophages. (E) *Efcab6*-expressing macrophages (marked by *Adgre1*) localize near a stage IV-VI seminiferous tubule, but not close to tubules at other stages. The basement membrane of the tubules are outlined by white dashed lines. Scale bar, 20 μm. (F) *Ghitm*/fm-expressing macrophages (marked by *Adgre1*) localize near a stage I-III seminiferous tubule, but not close to tubules at other stages. The basement membrane of the tubules are outlined by white dashed lines. Scale bar, 20 μm.

Similar to the Leydig cells, we performed differential gene expression analysis on macrophage populations (Fig 3D). We then used smFISH to confirm the stage-specific expression pattern of two of these genes *Efcab6* (Fig 3E, white arrow) and *Ghitm* (Fig 3F, white arrow). Moreover, recent studies have uncovered two populations of macrophages from two distinct lineages, with one population localized in the interstitial space and the other in the peritubular space (DeFalco et al., 2015; Mossadegh-Keller et al., 2017). We distinguished these two populations in the spatial transcriptome atlas based on their relative spatial position with Leydig cell beads since Leydig cells are known to occupy the interstitial space (Fig S6B). We found that beads assigned as peritubular macrophages exclusively expressed *H2-Ab1* and *Il1b* genes (Fig 3F), consistent with previous findings that these two genes are marker genes for peritubular macrophages (Mossadegh-Keller et al., 2017). Together, we demonstrated how the cycle of the seminiferous epithelium influenced the molecular composition of Leydig cells and macrophages by identifying the stage-dependent gene expression programs. Our analysis also successfully distinguished the two lineages of testicular macrophages based on their spatial localization.

### Building a Spatial Transcriptome Atlas for Human Spermatogenesis

Given the successful establishment of the mouse testicular spatial transcriptome atlas, we next used the same computational pipeline to spatially map human testicular cell types (Fig 4A & Fig S7A). The accuracy of the cell type assignment was validated by the enrichment of cell type marker genes (Fig S7B). Moreover, the structure of the reconstructed pseudotime image using the human Slide-seq data agreed with the morphology of the adjacent cross-section (Fig S8). Using the same method, we identified 788 human SP genes (Table S2), of which 580 genes showed matching spatial expression patterns with data from the Human Protein Atlas (https://www.proteinatlas.org/) (Uhlén et al., 2015). The rest of genes were either not covered in the database or showed non-specific immunostaining signals. For example, the human spatial transcriptome atlas identified SP genes *ACTRT3* and *ACRBP* to be expressed near the tubule lumen and genes *CLDN11* and *GSTA1* along the basement membrane. These observations were consistent with the corresponding protein distributions (Fig 4B). Next, we assigned stage information to each seminiferous tubule using genes with known stage-dependent patterns (Klaus et al., 2016), which resulted in the assignment of 3 stage clusters (stage I-II, III-IV, and V-VI). We found that unlike the mouse *Tnp1* gene which showed a stage-dependent spatial expression pattern (Fig 2D), the spatial expression pattern of human *TNP1* gene remained constant throughout different stages (Fig 4C). We also showed that *LPIN1*, a gene encoding a protein with phosphatide phosphatase and transcriptional coactivator activities (Reue, 2009) exhibited previously unidentified spatial expression patterns across stages which were confirmed by smFISH experiments (Fig S9A). Finally, GO enrichment analysis of SP genes showed both shared and stage-dependent signaling pathways among different stage clusters (Fig S9B). For example, SP genes regulating spermatid development and differentiation were shared by all stage clusters whereas SP genes associated with fertilization pathways such as *SMCP*, *TEX101*, and *AKAP3* were preferentially expressed in stage I-II tubules (Fig S9B).

**Figure 4.**
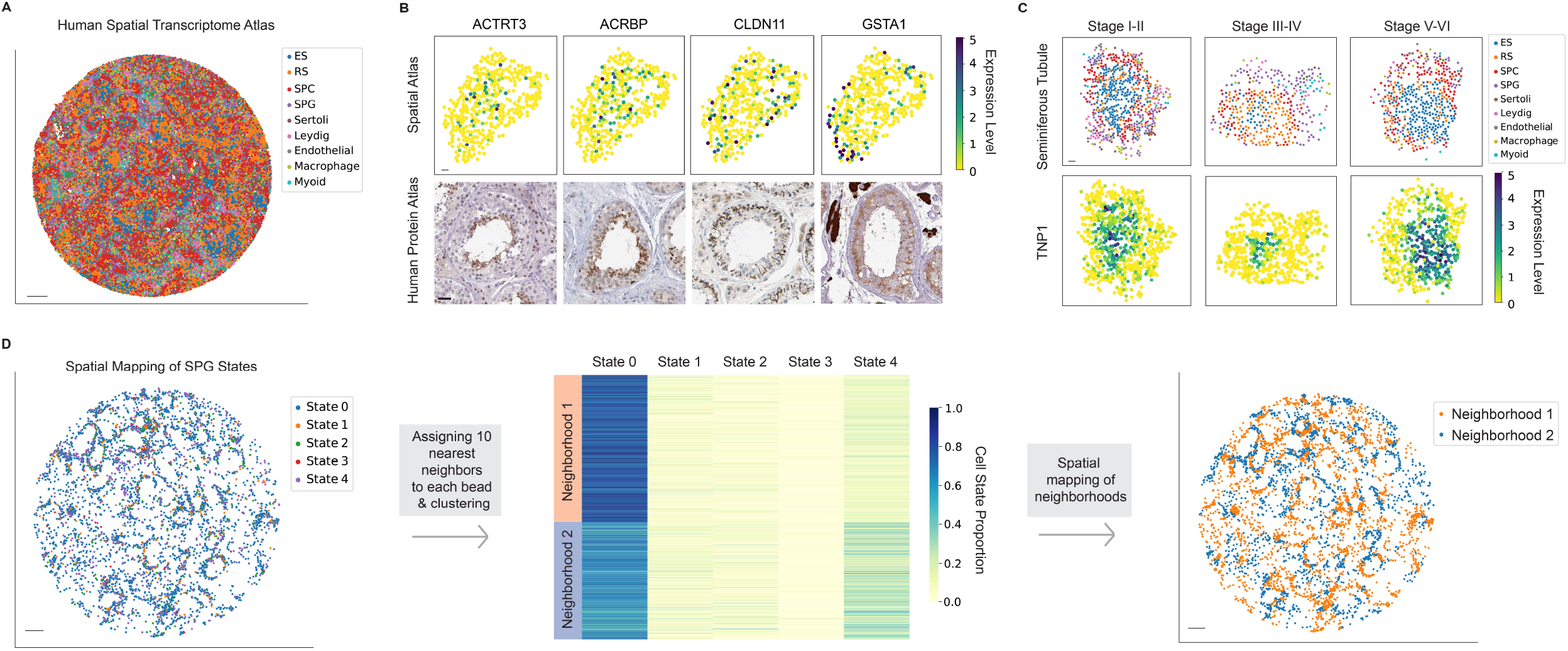
The Human Testicular Spatial Transcriptome Atlas. (A) Spatial mapping of testicular cell types. ES, elongating/elongated spermatid; RS, round spermatid; SPC, spermatocyte; SPG, spermatogonium. Scale bar, 300 μm. (B) The spatial expression patterns of genes *ACTRT3, ACRBP, CLDN11*, and *GSTA1* revealed by the human testicular spatial transcriptome atlas and the human protein atlas. Scale bar, 30 μm for the upper panel; 40 μm for the lower panel. (C) The spatial expression pattern of *TNP1* across stage clusters. Scale bar, 30 μm. (D) Five transcriptional states of the human spermatogonial population are spatially mapped (left). Two spatially segregated cellular neighborhoods are identified by clustering on cell state proportions (middle). Spatial distribution of the two neighborhoods (right). Scale bar, 300 μm.

### Spatial mapping of the transcriptional states of human spermatogonia

scRNA-seq has recently revealed extensive molecular heterogeneity among the human spermatogonium population. Five sequential transcriptional states (state 0-4) have been characterized among the entire human spermatogonia, with state 0 likely representing the earliest/naive spermatogonial stem cells (SSCs) (Guo et al., 2018). To spatially map these transcriptional states, we extracted beads assigned as spermatogonia in the human spatial atlas and performed cell state assignment to match the five transcriptional states (Fig 4D & Fig S10A). The accuracy of the state assignment was confirmed by the enrichment of state marker genes (Fig S10B). To further examine the spatial distribution of the five states, we performed a cellular neighborhood analysis (Methods). Two major cellular neighborhoods were identified, with neighborhood 1 containing mostly a subpopulation of state 0 SSC beads, and neighborhood 2 encompassing most of state 1-4 beads and the remaining state 0 SSC beads (Figure 4D). These two neighborhoods can be found within the same seminiferous tubule, but at distinct locations (Figure 4D), suggesting two different spermatogonial microenvironments composed of spermatogonia at distinct transcriptional states existing in the same seminiferous tubule.

### Characterizing changes in seminiferous tubule structure during diabetes-induced testicular injuries

Finally, we envisioned that the Slide-seq workflow and the accompanying computational pipelines could also be applied to the testicular samples from genetic models. To this end, we generated slide-seq data on six cross-sections from three leptin-deficient diabetic mice (*ob/ob*) and three cross-sections from three matching wild type (WT) mice (one representative dataset shown in Fig 5A). An important complication of diabetes mellitus (DM) is the disturbance in the male reproductive system. Numerous reports are available on impaired spermatogenesis under diabetic conditions (Alves et al., 2013; Ding et al., 2015; Jangir and Jain, 2014; Maresch et al., 2019) although the underlying mechanisms of which have not been fully appreciated. We first performed differential gene expression analysis to identify the differentially expressed genes (DE genes) between *ob/ob* and WT samples in different cell types (Table S3). We found that genes mainly expressed in ESs such as *Prm1, Prm2, Odf1*, and *Smcp* were significantly downregulated, consistent with the phenotype of ES/spermatozoon loss in *ob/ob* testes reported by a previous study (Bhat et al., 2006). In contrast, multiple mitochondrial genes such as *mt-Nd1*, *mt-Rnr1*, *mt-Rnr2* as well as non-coding RNA *Malat1* were among the genes whose expression elevated in *ob/ob* testes. This was also consistent with the observations that the increased expression levels of mitochondrial-encoded genes were associated with mtDNA damage and mitochondrial dysfunction in the pathogenicity of DM (Antonetti et al., 1995), and the expression of *Malat1* was elevated in multiple diabetic-related diseases (Abdulle et al., 2019). In addition, we also identified genes involved in meiosis were differentially expressed such as *Ubb, Mael* and *Hspa2*, which echoed the findings in the ovary where the meiotic regulation in oocytes was altered in diabetic mice and rats (Colton et al., 2002; Kim et al., 2007). Together, these results demonstrated that the diabetic spatial atlas faithfully captured the molecular changes in diabetic testes.

**Figure 5.**
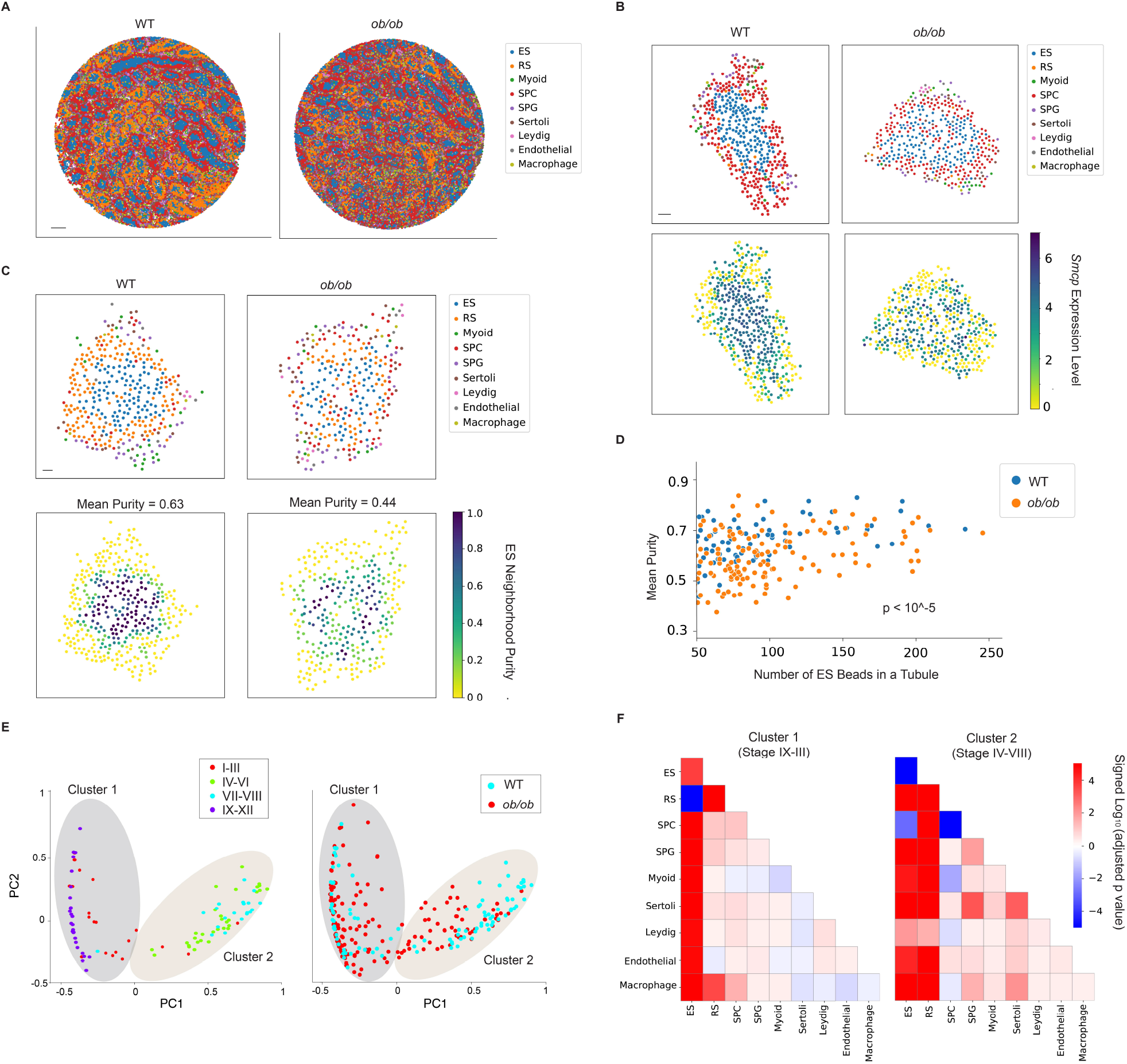
Diabetes causes disruptions in spatial arrangements of testicular cell types in the seminiferous tubules. (A) Spatial mapping of testicular cell types for wild type (WT) and ob/ob samples. Scale bar, 500 μm. (B) The spatial expression pattern of *Smcp* is disrupted in a representative ob/ob seminiferous tubule. ES, elongating/elongated spermatid; RS, round spermatid; SPC, spermatocyte; SPG, spermatogonium. Scale bar, 30 μm. (C) The ES purity score for each Slide-seq bead in a representative WT and ob/ob seminiferous tubule, respectively. The mean purity score for the beads with non-zero value in each tubule is also shown. Scale bar, 30 μm. (D) The mean purity score for each seminiferous tubule with at least 50 ES beads from the WT and ob/ob samples. Comparison of the purity score between the two conditions is performed using the Mann-Whitney U test. (E) Left: Two clusters of WT seminiferous tubule structures revealed by the principal components of pairwise spatial contact frequencies in WT seminiferous tubules. The stage information of each tubule is labeled. Right: assignment of ob/ob seminiferous tubules into the two clusters shown in the left plot. (F) Mann-Whitney U test on pairwise spatial cellular contact frequencies between WT and ob/ob seminiferous tubules under the two clusters. Signed p value of significant increases (positive) and decreases (negative) in spatial contact frequencies between cell types are shown.

Next, we sought to spatially map the expressions of the identified DE genes. Of interest, we noticed that some of the DE genes such as *Smcp, Odf1*, and *Malat1* showed altered spatial expression patterns in ob/ob testis cross-sections (Fig S11). By zooming into individual seminiferous tubules, we found that the spatial expression pattern of *Smcp* in WT testes was disrupted in *ob/ob* seminiferous tubules (Fig 5B). Since we have shown that the spatial expression pattern of *Smcp* was not stage-dependent (Table S1), such a change in the *ob/ob* tubules was not likely due to differences in stages of the seminiferous epithelium cycle. A close examination of the *ob/ob* tubules indicated that the change in gene expression pattern was in part, if not entirely due to an alteration in the spatial cellular organization especially among the spermatid population. To systematically and quantitatively profile such changes in *ob/ob* seminiferous tubules, we first devised a metric termed purity score to evaluate the extent of the spatial mixing between the ES beads with beads of other cell types (Methods). In a WT seminiferous tube, ES beads clustered together at the center, showing a high average purity score (Fig 5C, left). In contrast, such spatial organization was disrupted in *ob/ob* seminiferous tubules where ES beads were more likely to make spatial contacts with beads of other cell types, showing a low purity score (Fig 5C, right). We then calculated the average purity scores for 72 WT tubules and 136 *ob/ob* tubules and found that the scores were significantly lower in the *ob/ob* tubules than in the WT tubules (Fig 5D, Mann-Whitney U test). Next, we quantified the spatial arrangements of all the cell types within a seminiferous tubule by calculating the pairwise spatial contact frequencies (Methods). t-SNE (t-distributed stochastic neighbor embedding) analysis of the WT seminiferous tubules based on the pairwise spatial contact frequencies revealed two major clusters (Fig 5E, left). The separation of these two clusters was mainly contributed by differences in cell type proportions under different stages of the seminiferous epithelium cycle (Fig 5E, left). We then clustered the *ob/ob* seminiferous tubules in the same t-SNE space with the WT tubules (Fig 5E, right). By comparing each pairwise spatial contact frequency between the WT and *ob/ob* seminiferous tubules under the same cluster (Mann-Whitney U test), we effectively eliminated the influence of the tubule stages on cell type proportions. Consistent with the purity score measurement, we observed significant changes in the spatial arrangements between ES beads and beads of all other cell types (Fig 5F). Moreover, the spatial contacts between RS and other cell types, especially macrophages, were also markedly enhanced in *ob/ob* tubules (Fig 5F), suggesting that the spatial arrangement of spermatids were the most susceptible to diabetic conditions. Taken together, our analysis showed that the disruption in the spatial structure of seminiferous tubules was prevalent in diabetes-induced testicular injuries.

## Discussion

Spermatogenesis takes place in the spatially-confined environment of seminiferous tubules and is constantly influenced by the somatic cell types in the peritubular and interstitial space. However, there has been a lack of tools to comprehensively profile spermatogenesis within the native context of seminiferous tubules as well as to capture the spatial interactions between somatic cells and germ cells at both the molecular and cellular level. Moreover, within any given testis cross-section, each seminiferous tubule is at a specific stage with each stage associated with unique combinations of germ cell subtypes and biological events, making it challenging to obtain stage-specific spermatogenesis with molecular resolution. Using Slide-seq, we have generated an unbiased spatial transcriptome atlas for the mouse and human spermatogenesis at near single-cell resolution. By applying a custom computational pipeline, we were able to assign information of cell type, seminiferous tubule, and stage to each Slide-seq bead with high accuracy. Our data provides a novel framework to systematically evaluate the spatial dynamics in gene expressions and in cellular structures both within and surrounding the seminiferous tubules with high resolution.

### The Spatial Transcriptome Atlas as a Platform for Profiling Spatial Gene Expression Dynamics in Spermatogenesis

The testis expresses the largest number of genes of any mammalian organ and many of these genes are testis-specific (Brawand et al., 2011; Djureinovic et al., 2014; Mele et al., 2015). However, the localization and biological functions of these genes remain largely unexplored. Here, we showed the spatial transcriptome atlas as a powerful platform for profiling spatial gene expression dynamics: 1) using the spatial atlas, we systemically localized genes at the level of individual seminiferous tubules in both the mouse and human testis; 2) we observed changes in the spatial expression pattern of the same gene under different stages of the seminiferous epithelium cycle; 3) due to the cyclic nature of spermatogenesis and the availability of the stage information, we were able to use the spatial transcriptomic data to infer temporal expression dynamics of the genes along the developmental trajectory of germ cells. This provided information on the germ cell developmental stages during which these genes are likely to exert their functions; 4) by grouping the spatially patterned genes (SP genes) based on their involvements in different signaling pathways, we nominated genes with previously underappreciated functions; and 5) besides SP genes, we used the spatial transcriptome atlas to also identify genes with stage-specific expression in two somatic cell types of the testis: Leydig cells and macrophages. These genes may exert their functions in a stage-dependent manner.

### The Spatial Transcriptome Atlas as a Platform to Profile Spatial Cellular Structures of Seminiferous Tubules

In both the mouse and human testis, testicular cell types are spatially organized in a stereotypic pattern within and surrounding the seminiferous tubules to support spermatogenesis. A comprehensive profiling of these spatial cellular structures under both homeostasis and pathology would provide useful information on spermatogenesis and disease progression. In this study, we spatially mapped human subpopulations of spermatogonia at different transcriptional states in the seminiferous tubules. A cellular neighborhood identification analysis identified two spatially segregated neighborhoods, with the first neighborhood mainly composed of a subpopulation of the spermatogonial stem cells and the second made up of the rest of the stem cells and cells from other transcriptional states. The existence of the two neighborhoods in the same seminiferous tubule may suggest two distinct spermatogonial stem cell niches in human seminiferous tubules. In fact, a similar observation has been made in the mouse seminiferous tubules that outside of the niche where A_Single_/A_Paired_/A_Aligned_ spermatogonia are located, there exists a separate niche structure harboring the so-called ‘ultimate stem cells’ which are high in *Id4* expression (de Rooij, 2017). Although it is tempting to draw parallels between these two models, more evidence is needed to pinpoint the proposed two niche structures in the human seminiferous tubules.

In addition to profiling normal testis samples, we also systematically compared the spatial cellular structures in WT and diabetic seminiferous tubules. By calculating the extent of spatial mixing and pairwise cell-cell contact frequencies between different cell types, we observed a significant disruption in spatial cellular organization of the diabetic seminiferous tubules. Of interest, a previous study using second order stereology has indicated that the spatial arrangement of Sertoli cells and spermatogonia is significantly disrupted in the diabetic testes (Sajadi et al., 2019). Indeed, our analysis showed a significant change in the spatial contact frequencies between Sertoli cells and spermatogonia. Taken together, a disruption in the spatial cellular organization at the level of seminiferous tubules is a prevalent phenotype in diabetes-induced testicular injuries. Although we only applied Slide-seq to the diabetic models, we envision that it can be readily adapted to profile the spatial cellular structures in other perturbation models.

In summary, the mouse and human testicular spatial transcriptome atlas are valuable resources to comprehensively reveal the detailed spatial molecular and cellular information that instructs spermatogenesis. The ability to profile systematically, quantitatively, and in a spatially resolved manner the genome-wide changes of gene expression as well as the spatial cellular organization at near singlecell resolution has a transformative potential in the field of reproductive biology. The Slide-seq protocol and analytical framework can be readily adapted to other stereotypically structured tissues and developing animals and might be applicable to more perturbation models as well.

### Contact for Reagent and Resource Sharing

Further information and requests for resources and reagents should be directed to and will be fulfilled by the Lead Contact, Fei Chen.

### Experimental Model and Subject Details

All animal experiments were carried out with prior approval of the Broad Institute of MIT and Harvard on Use and Care of Animals (Protocol ID: 0211-06-18), in accordance with the guidelines established by the National Research Council Guide for the Care and Use of Laboratory Animals. Adult (7 to 18 weeks old) male mice were housed in the Broad Institute animal facility, in an environment controlled for light (12 hours on/off) and temperature (21 to 23 °C) with *ad libitum* access to water and food. For detailed mouse strain information, see Key Resources Table.

Adult human testicular samples for Slide-seq and smFISH were from two healthy men (donor #1: 25 years old; donor #2: 32 years old); Samples were obtained through the University of Utah Andrology laboratory and Intermountain Donor Service. Both samples were de-identified.

## Method Details

### Transcardial Perfusion

Mice were anesthetized by administration of isoflurane in a gas chamber flowing 3% isoflurane for 1 minute. Anesthesia was confirmed by checking for a negative tail pinch response. Animals were moved to a dissection tray and anesthesia was prolonged via a nose cone flowing 3% isoflurane for the duration of the procedure. Transcardial perfusions were performed with ice cold pH 7.4 PBS to remove blood from the testes. Testes were removed and frozen for 3 minutes in liquid nitrogen vapor and moved to −80°C for long term storage.

### Slide-seq workflow

The workflow of Slide-seq was described previously (Stickels et al., 2020). Briefly, the 10-μm polyT barcoded beads were synthesized in house. Each bead oligo contains a linker sequence, a spatial barcode, a UMI sequence, and a poly(dT) tail. The bead arrays were prepared by pipetting the synthesized beads which were pelleted and resuspended in water + 10% DMSO at a concentration between 20,000 and 50,000 beads/μL, into each position on a gasket. The coverslip-gasket filled with beads was then centrifuged at 40°C, 850g for at least 30 minutes until the surface was dry. In situ sequencing of the bead array to extract the spatial barcodes on the beads were performed in a Bioptechs FCS2 flow cell using a RP-1 peristaltic pump (Rainin), and a modular valve positioner (Hamilton MVP). Flow rates between 1 mL/min and 3 mL/min were used during sequencing. Imaging was performed using a Nikon Eclipse Ti microscope with a Yokogawa CSU-W1 confocal scanner unit and an Andor Zyla 4.2 Plus camera. Images were acquired using a Nikon Plan Apo 10x/0.45 objective. Sequencing was performed using a sequencing-by-ligation approach. Image processing was performed using a custom Matlab package (https://github.com/MacoskoLab/PuckCaller/).

Fresh frozen testis tissue was warmed to −20°C in a cryostat (Leica CM3050S) for 20 minutes prior to handling. Tissue was then mounted onto a cutting block with OCT and sliced at 10 μm thickness. Sequenced bead arrays were then placed on the cutting stage and tissue was maneuvered onto the pucks. The tissue was then melted onto the array by moving the array off the stage and placing a finger on the bottom side of the glass. The array was then removed from the cryostat and placed into a 1.5 mL eppendorf tube. The sample library was then prepared as below. The remaining tissue was re-deposited at −80 °C and stored for processing at a later date.

Bead arrays with tissue sections were immediately immersed in 200 μL of hybridization buffer (6x SSC with 2 U/μL Lucigen NxGen RNAse inhibitor) for 30 minutes at room temperature to allow for binding of the mRNA to the oligos on the beads. Subsequently, first strand synthesis was performed by incubating the pucks in 200 μL of reverse transcription solution (Maxima 1x RT Buffer, 1 mM dNTPs, 2 U/μL Lucigen NxGen RNAse inhibitor, 2.5 μM template switch oligo with Qiagen #339414YCO0076714, 10 U/μL Maxima H minus reverse transcriptase) for 1.5 hours at 52 °C. 200 μL of 2x tissue digestion buffer (200 mM Tris-Cl pH 8, 400 mM NaCl, 4% SDS, 10 mM EDTA, 32 U/mL Proteinase K) was then added directly to the reverse transcription solution and the mixture was incubated at 37°C for 30 minutes. The solution was then pipetted up and down vigorously to remove beads from the glass surface, and the glass substrate was removed from the tube using forceps and discarded. 200 μL of wash buffer (10 mM Tris pH 8.0, 1 mM EDTA, 0.01% Tween-20) was then added to the solution mix and the tube was then centrifuged for 3 minutes at 3000 RCF. The supernatant was then removed from the bead pellet, the beads were resuspended in 200 μL of wash buffer and were centrifuged again. This was repeated a total of three times. The supernatant was then removed from the pellet. The beads were then resuspended in 200μl of 1 U/μL Exonuclease I and incubated at 37 °C for 50 minutes. After Exonuclease I treatment the beads were centrifuged for 3 minutes at 3000 RCF. The supernatant was then removed from the bead pellet, the beads were resuspended in 200 μL of wash buffer and were centrifuged again. This was repeated a total of three times. The supernatant was then removed from the pellet. The pellet was then resuspended in 200 μl of 0.1 N NaOH and incubated for 5 minutes at room temp. To quench the reaction 200 μl of wash buffer was added and beads were centrifuged for 3 minutes at 3000 RCF. The supernatant was then removed from the bead pellet, the beads were resuspended in 200 μL of wash buffer and were centrifuged again. This was repeated a total of three times. Second strand synthesis was then performed on the beads by incubating the pellet in 200μl of second strand mix (Maxima 1x RT Buffer, 1 mM dNTPs, 10 μm dN-SMRT oligo, 0.125 U/μL Klenow fragment) at 37 °C for 1 hour. After the second strand synthesis 200μl of wash buffer was added and the beads were centrifuged for 3 minutes at 3000 RCF. The supernatant was then removed from the bead pellet, and the beads were resuspended in 200 μL of wash buffer and were centrifuged again. This was repeated a total of three times. 200 μl of water was then added to the bead pellet and the beads were moved into a 200 μL PCR strip tube, pelleted in a mini-centrifuge, and resuspended in 200 μL of water. The beads were then pelleted and resuspended in PCR mix (1x Terra Direct PCR mix buffer, 0.25 U/μL Terra polymerase, 2 μM Truseq PCR handle primer, 2 μM SMART PCR primer) and PCR was performed using the following program: 95 °C 3 minutes; 4 cycles of: 98 °C for 20 s, 65 °C for 45 s, 72 °C for 3 min and 9 cycles of: 98 °C for 20 s, 67 °C for 20 s, 72 °C for 3 min; 72 °C for 5 min and hold at 4 °C. The PCR product was then purified using 30 μL of Ampure XP beads (Beckman) according to manufacturer’s instructions and resuspended into 50 μL of water and the cleanup was repeated and resuspended in a final volume of 10 μl. 1 μL of the library was quantified on an Agilent Bioanalyzer High sensitivity DNA chip (Agilent). Then, 600 pg of PCR product was prepared into Illumina sequencing libraries through tagmentation with a Nextera XT kit (Illumina). Tagmentation was performed according to manufacturer’s instructions and the library was amplified with primers Truseq5 and N700 series barcoded index primers. The PCR program was as follows: 72°C for 3 min and 95°C for 30 s; 12 cycles of: 95°C for 10 s, 55°C for 30 s, 72°C for 30 s; 72°C for 5 min and hold at 10°C. Samples were cleaned with AMPURE XP (Beckman Coulter A63880) beads in accordance with manufacturer’s instructions at a 0.6x bead/sample ratio (30 μL of beads to 50 μL of sample) and resuspended in 10μL of water. Library quantification was performed using the Bioanalyzer. Finally, the library concentration was normalized to 4 nM for sequencing. Samples were sequenced on the Illumina NovaSeq S2 flow cell 100 cycle kit with 12 samples per run (6 samples per lane) with the read structure 42 bases Read 1, 8 bases i7 index read, 50 bases Read 2. Each bead array received approximately 150-200 million reads, corresponding to ~2,000-2,500 reads per bead.

### Immunofluorescence

Slides with tissue sections were fixed in 4% PFA (in PBS) and washed three times in PBS for 5 minutes each, blocked 30 minutes in 4% BSA (in PBST), incubated with primary antibodies overnight at 4°C, washed three times in PBS for 5 minutes each, and then incubated with secondary antibodies at for 1 hour at room temperature. Slides were then washed twice in PBS for 5 minutes each and then for 10 minutes with a PBS containing DAPI (D9542, Sigma-Aldrich). Lastly, slides were mounted using ProLong™ Gold Antifade Mountant (P36934, Thermo Fisher Scientific) and sealed. Antibodies used for IF: Rabbit Anti-HABP4 antibody (1:100, HPA055969, Millipore Sigma), mouse anti-phospho-histone H2A.X (Ser139) antibody, FITC conjugate (1:100, 16-202A, Millipore Sigma), Sheep anti-acetyl histone H4 antibody (1:100, AF5215, R&D Systems), and Alexa Fluor 594-, and 647-conjugated secondary antibodies (Thermo Fisher Scientific) were used.

### Validation by Single Molecule RNA HCR

Single molecule RNA HCR was performed as described by Choi et al (Choi et al., 2018) with small modifications. Briefly, frozen testes were sectioned into (3-Aminopropyl) triethoxysilane-treated glass bottom 24-well plates. Sections were cross-linked with 10% Formalin for 15 min, washed with 1X PBS for three times, and permeabilized in 70% ethanol for 2 hours overnight at −20 °C. Sections were then rehydrated with 2X SSC (ThermoFisher Scientific) for three times and equilibrated in HCR hybridization buffer (Molecular Instruments, Inc.) for 10 minutes at 37 °C. Gene probes were added to the sections at a final concentration of 2 nM in HCR hybridization buffer and hybridized overnight in a humidified chamber at 37 °C. Custom probe sets were designed and synthesized by Molecular Instruments, Inc.. After hybridization, samples were washed four times in HCR wash buffer (Molecular Instruments, Inc.) for 15min at 37 °C and then three times for 5 minutes at room temperature in 5X SSCT (ThermoFisher Scientific). The probe sets were amplified with HCR hairpins for 12-16 hours at room temperature in HCR amplification buffer (Molecular Instruments, Inc.). Fluorescently-conjugated DNA hairpins used in the amplification were ordered from Molecular Instruments, Inc.. Prior to use, the hairpins were ‘snap cooled’ by heating at 95 °C for 90 seconds and letting cool to room temperature for 30 min in the dark. After amplification, the samples were washed in 5X SSCT and stained with 20ng/mL DAPI (Sigma) before imaging. Microscopy was performed using an inverted Nikon CSU-W1 Yokogawa spinning disk confocal microscope with 488 nm, 640nm, 561nm, and 405 nm lasers, a Nikon CFI APO LWD 40X/1.15 water immersion objective, and an Andor Zyla sCMOS camera. NIS-Elements AR software (v4.30.01, Nikon) was used for image capture.

### Computational Methods for Slide-seq Data

#### Preprocessing of Slide-seq Data

The Slide-seq tools (https://github.com/MacoskoLab/slideseq-tools) were used for processing raw sequencing data. In brief, the Slide-seq tools extracted the barcode for each read in an Illumina lane from Illumina BCL files, collected, demultiplexed, and sorted reads across all of the tiles of a lane via barcode, and produced an unmapped bam file for each lane. The bam file was tagged with cellular barcode (bead barcode) and molecular barcode (UMI). Low-quality reads were filtered, and reads were trimmed with starting sequence and adapter-aware poly A. STAR (Dobin et al., 2013) was used to align reads to genome sequence. The unmapped bam and sorted aligned bam were merged and tagged with interval and gene function. For each of those barcodes with short read sequencing, hamming distances were calculated between it and all of the bead barcodes from in situ sequencing. The list of unique matched Illumina barcodes with hamming distance <= 1 along with the matched bead barcodes were outputted. The final output from the Slide-seq tools were a digital gene expression matrix (bead barcodes X genes) and barcode location matrix (bead barcodes X spatial coordinates). For each Slide-seq bead, the total number of UMIs were calculated and beads with less than 20 UMIs were filtered out. A trimming step was also applied to exclude beads located outside the main bead array area.

#### Cell Type Assignment

To accurately assign cell type to each Slide-seq bead, the contribution of each cell type to the RNA on the bead was computed using a custom method, termed NMFreg (Non-Negative Matrix Factorization Regression) (Rodriques et al., 2019). The method consisted of two main steps: first, single-cell atlas data previously annotated with cell type identities was used to derive a basis in reduced gene space (via NMF), and second, non-negative least squares (NNLS) regression was used to compute the loadings for each bead in that basis. To perform NMF on the mouse slide-seq data, we used scRNA-seq data from a study by Green et al (Green et al., 2018). For the human, we used data from Guo et al. as a reference (Guo et al., 2018). The cell type of the bead was assigned based on the identity of the maximum factor loading determined by NMFreg.

#### Pseudotime Reconstruction

To assign pseudotime value to each Slide-seq bead, first we took a testis scRNA-seq dataset as an input and assigned a pseudotime value to each cell from that dataset using Monocle (Qiu et al., 2017). For the mouse, we used a published scRNA-seq dataset from Lukassen et al (Lukassen et al., 2018). For the human, we used the data from Guo et al. (Guo et al., 2018). Next, we used Monocle to identify 1000 genes whose expression changed as a function of pseudotime. We then selected genes whose expression co-varied the most with pseudotime using L1 regularization. For the mouse data, we iterated the feature selection process through 1-20 genes and selected 14 genes which minimized the number of genes to be used while maximizing the fidelity of pseudotime reconstruction. The resulting linear function returned by L1 regularization included 6 genes with negative weights (hereafter referred as negative genes) and 8 genes with positive weights (positive genes). The negative genes and the corresponding coefficients are: *mt-Nd1*: 2.240772116330779, *Tuba3b*: 20.0, *Stmn1*: 4.661355144844998, *Cypt4*: 2.914946084728282, *mt-Cytb*: 7.940721195057547, *Hsp90aa1*: 7.719926741335844; The positive genes and the corresponding coefficients are *Tnp2*: 2.2300113564016115, *Smcp*: 20.0, *Gsg1*: 10.749147113890643, *Oaz3*: 13.684470608169942, *Hmgb4*: 11.19717467780924, *Lyar*: 3.205774497366639, *Prm1*: 1.8021648809899742, *Dbil5*: 2.4081140269931445. For the human, we selected 11 negative genes and 6 genes with positive genes. The negative genes and the corresponding coefficients are *TNP1*: 0.9106007166752985, *RPL39*: 9.212049749934977, *PTGDS*: 5.771773680502793, *RPLP2*: 20.0, *FTH1*: 19.992459791415897, *MALAT1*: 17.76205459561361, *DCN*: 19.475013103219712, *RPLP1*: 18.860848405765406, *CFD*: 7.067158789040227, *FTL*: 16.55433814467757; The positive genes and their corresponding coefficients are *TSACC*: 1.3896196907421736, *NUPR2*: 20.0, *TMSB4X*: 3.638712833769793, *PRM2*: 1.2032600855690998, *PRM1*: 0.4449200139269451. We then extracted the UMI count for each gene in the gene list from the Slide-seq data and multiplied that with the corresponding weight. The pseudotime value for each slide-seq bead was calculated by summing the weighted UMI count value from every gene in the gene list.

#### Segmentation of Seminiferous Tubules

The fact that the image of pseudotime reconstruction retains the morphological structure of seminiferous tubules suggests that the image can be used to extract tubule information. Each slide-seq bead can be treated as a pixel and the pseudotime value assigned to each bead can be viewed as pixel value. To this end, we developed a custom computational pipeline to first convert the pseudotime reconstruction into a grayscale image. Next, the grayscale image was smoothed using Gaussian blur with a sigma of 2. The smoothed image was then thresholded with an arbitrary pixel value cutoff which isolated individual tubes while retaining a maximum number of pixels passing the threshold. This was followed by a standard watershed workflow to find local maxima, perform distance transform and segment individual seminiferous tubules. Finally, we performed *K*-nearest neighbor analysis to assign beads which were excluded during the thresholding step back to the nearest tubules. The segmented tubules were plotted and visually inspected. Wrongly assigned beads (for example, ES beads from the adjacent seminiferous tubules but assigned to the periphery of the current tubule) and beads with low cell type certainty as determined in the NMFreg step were excluded for subsequent analysis.

#### Assignment of stages

Segmented seminiferous tubules with a total number of beads less than 20 and more than 700 were first filtered out. Next, the raw UMI count data was normalized and variance-stabilized using the function SCTransform for Seurat V3 (Hafemeister and Satija, 2019). The normalized gene expression values of the Slide-seq beads within the same tubule were aggregated. Uniform manifold approximation and projection (UMAP) for dimension reduction of the aggregated gene expression data were performed using the top 3000 highly variable genes (HVGs) under the assumption that seminiferous tubules at similar stages should share similar transcriptional profiles. Finally, genes whose expressions are known to be stage-specific (Johnston et al., 2008; Klaus et al., 2016) were used to assign stages to each cluster.

#### GO analysis

For each SP gene group identified by spatial profiling, GO analysis was performed using the clusterProfiler package (Yu et al., 2012). Cellular components from the org.Mm.eg.db genome wide annotation for the mouse and org.Hs.eg.db for the human was used for the ontology database.

#### Assignment of interstitial and peritubular macrophages

A pairwise distance matrix was first calculated between macrophages and Leydig cell beads. Next, all macrophage beads were ranked based on their distances to the nearest Leydig cell beads. Macrophage beads with a distance less than 30 μm were assigned as interstitial, and the rest of the macrophage beads were peritubular.

#### Cellular neighborhood identification for the spermatogonial population

The neighborhood identification analysis was adapted from Schürch *et.al*. (Schürch et al., 2020). Briefly, we generated sliding windows with a window size of 10 (i.e., 10 beads per window) based on the spatial coordinates of the spermatogonium beads and their assigned state identities. For each window, we counted the number of each cell state that was present and calculated the frequency of each cell state within each window. Next, we performed unsupervised *K*-means clustering on the cell state frequency lists, effectively clustering the windows based on cell state frequency given a specific number of clusters, or neighborhoods. We tested different values for clusters/neighborhoods (i.e., *K*) and determined that K=2 lead to the most biologically relevant result.

#### Differential gene expression analysis between WT and *ob/ob* Slide-seq data

To enable directly comparative analyses within cell types between WT and ob/ob samples, we used Seurat 3 (v3.1) (Stuart et al., 2019) to perform joint analysis. The top 2,000 highly HVGs were identified using the function FindVariableFeatures with the vst method. CCA was used to identify common sources of variation between WT and *ob/ob* cells. The first 20 dimensions of the CCA were chosen to integrate the Slide-seq datasets from the two conditions. After integration, the expression level of HVGs in the cells was scaled and centered for each gene across cells, and PCA analysis was performed on the scaled data. The DE genes for each cell type was identified using the function FindMarkers.

#### Calculation of the purity score

To quantify the extent of spatial mixing between ES beads and beads of other cell types, we calculated the purity score of the ES bead neighborhoods. In short, a *K*-nearest neighbor was performed for each ES bead in a seminiferous tubule (*K* = 5 in this case). The percentage of the ES bead in each neighborhood (i.e., the purity of the neighborhood) was calculated. The purity scores for all the ES beads in the tubule were averaged to determine the mean purity score. Only tubules with at least 50 ES beads were considered.

#### Calculation of the pairwise spatial contact frequency

For each bead in a seminiferous tubule, we counted the number of edges (i.e., spatial contacts) between that bead and its 10 nearest neighboring beads whose cell types were also recorded. The spatial contacts between two cell types were normalized by the total number of spatial contacts among all the beads. A total of 45 pairwise spatial contact frequencies were calculated for each seminiferous tubule and were used for the principle component analysis and t-SNE projection.

#### Data Availability

The raw sequencing data supporting the findings of this study are available in NCBI BioProject database with BioProject ID PRJNA668433. The processed data including the gene expression matrices, bead location matrices, and the NMFreg-enabled cell type assignment matrices are available at https://www.dropbox.com/s/ygzpj0d0oh67br0/Testis_Slideseq_Data.zip?dl=0.

#### Code Availability

Custom code is available at https://github.com/thechenlab/Testis_Slide-seq.

## Acknowledgments

We thank Jamie Marshall for the help on mouse testicular sample collections; Dylan Cable, Aleks Goeva, Sarah Mangiameli, Dan Lesman, Sophia Liu and Yu Jiang for discussions on the computational analysis; Tongtong Zhao for comments on the project. F.C. acknowledges support from Eric and Wendy Schmidt as funders of the Schmidt Fellows Program at the Broad Institute. H.C. thanks the Lalor Foundation and the Male Contraceptive Initiative for the support.

## Author Contributions

F.C. supervised the project. H.C. conceived the project. H.C. and F.C. designed the experiments. H.C. and E.M. performed experiments. H.C., F.C., A.L. and J.L. performed data analysis. X.N., J.H., B.R.C. and J.G. provided human testicular samples. C.Y.C. and E.Z.M. provided consultations. H.C. and F.C. wrote the manuscript with input from all authors.

