## Supplemental Figures for "Dissecting Mammalian Spermatogenesis Using Spatial Transcriptomics": Supplemental Figures.pdf

### Figure S1

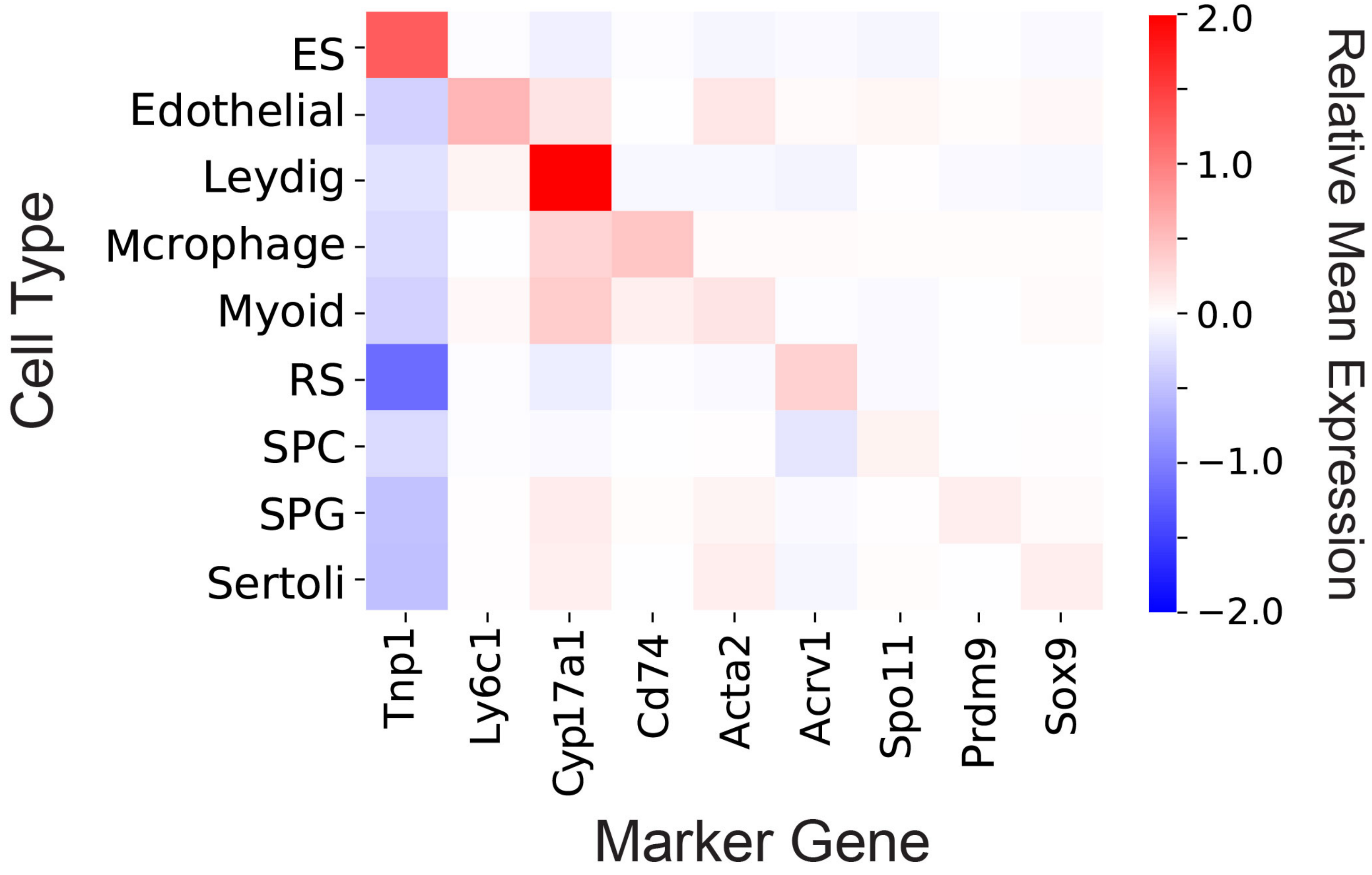


**Figure S1 Validation of the mouse testicular cell type assignment using marker genes**
The mean gene expression matrix for cell type marker genes. ES, elongating/elongated spermatid; RS, round spermatid; SPC, spermatocyte; SPG, spermatogonium.

### Figure S2

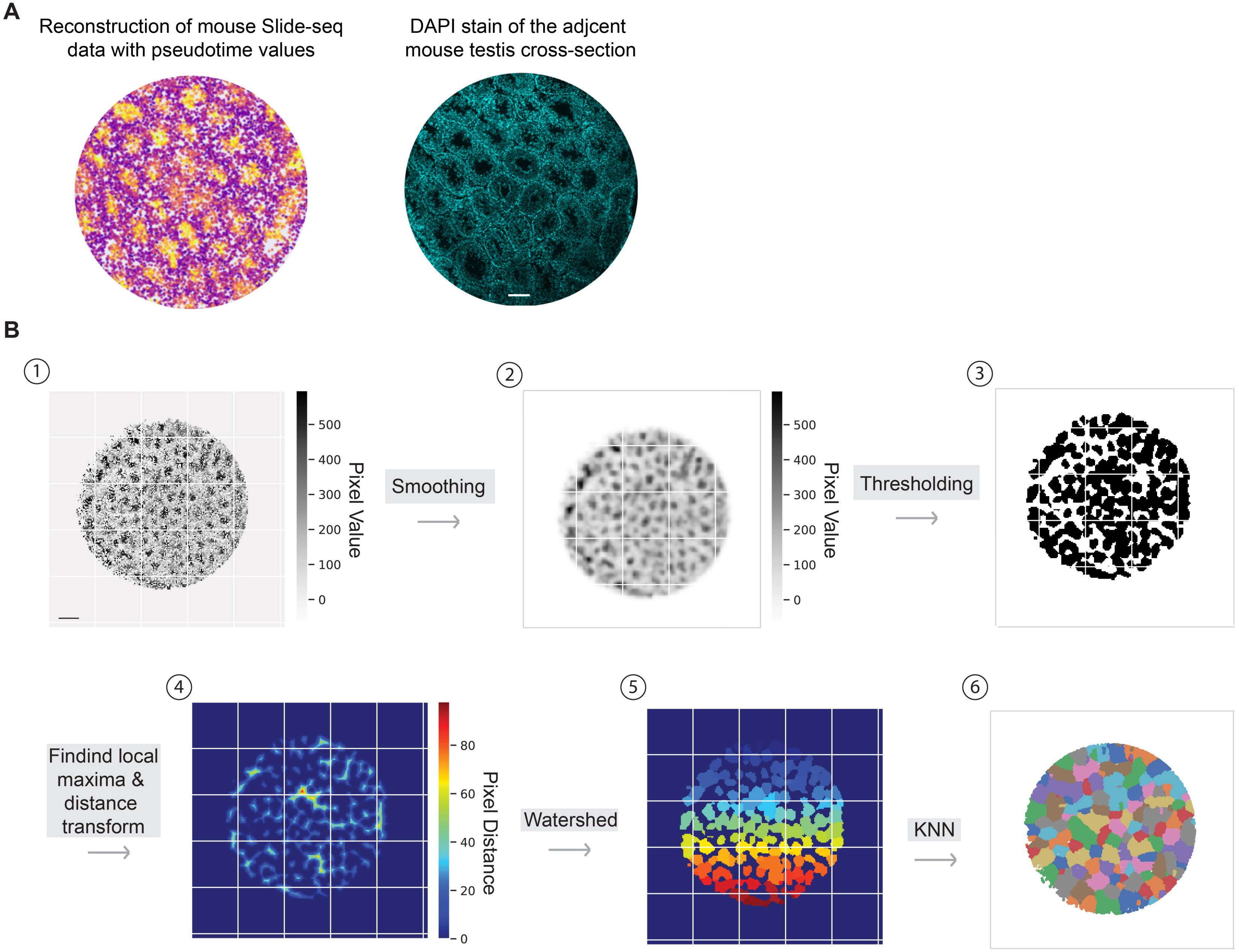


**Figure S2 Digital segmentation of the seminiferous tubules**

(A) The pseudotime-reconstructed image of the mouse Slide-seq data shows consistent morphological features with the DAPI stain image of the adjacent testis cross-section.Scale bar, 100 μm.

(B) Workflow of the segmentation on Slide-seq beads. Scale bar, 300 μm.

### Figure S3

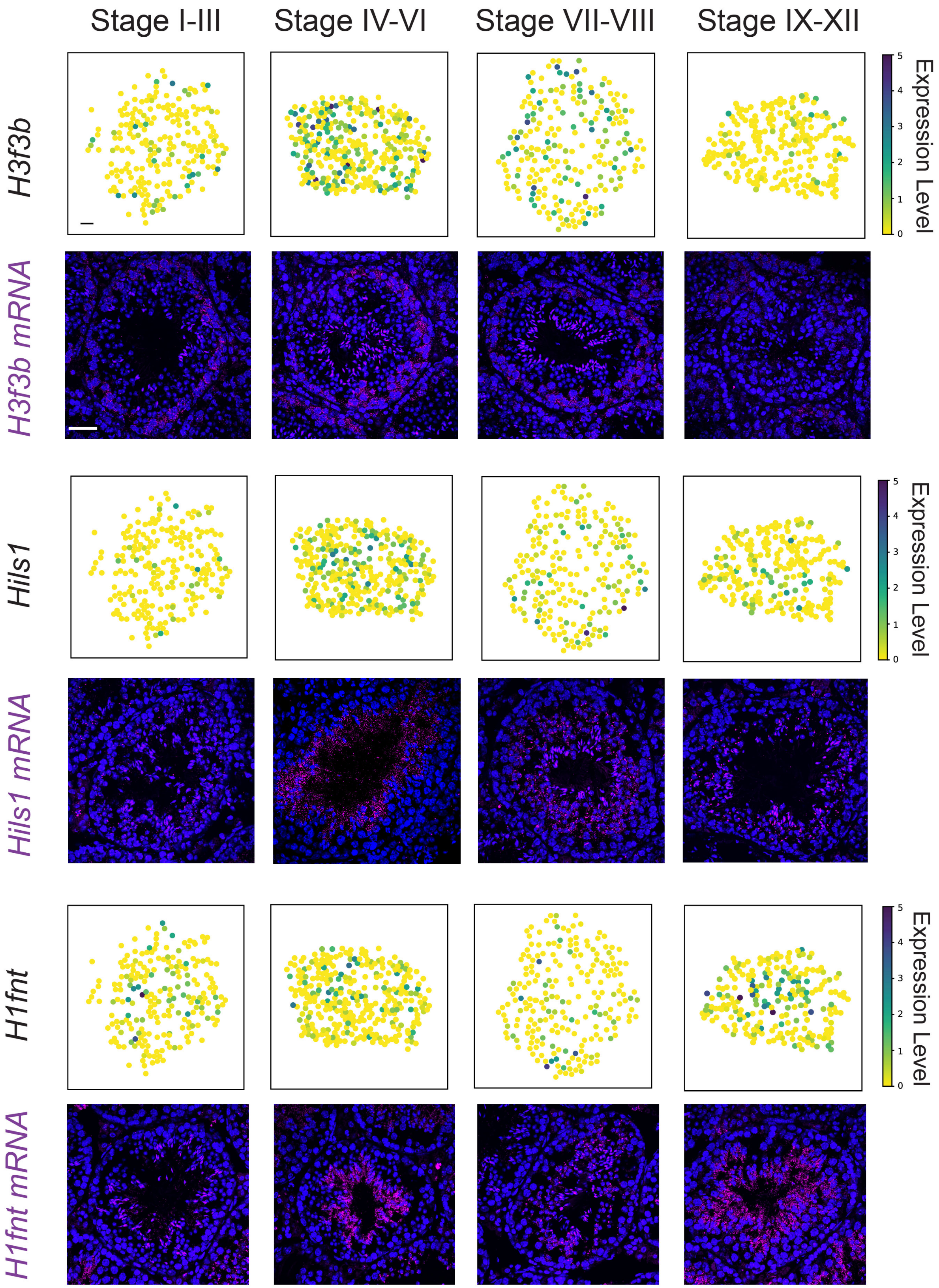


**Figure S3 Spatial gene expression patterns of histone components**
The spatial expression pattern of mouse histone genes *H3f3b*, *Hils1*, and *H1fnt* revealed by the spatial transcriptome atlas and smFISH. Scale bar, 30 μm for both the digitally reconstructed seminiferous tubule images and the smFISH images.

### Figure S4

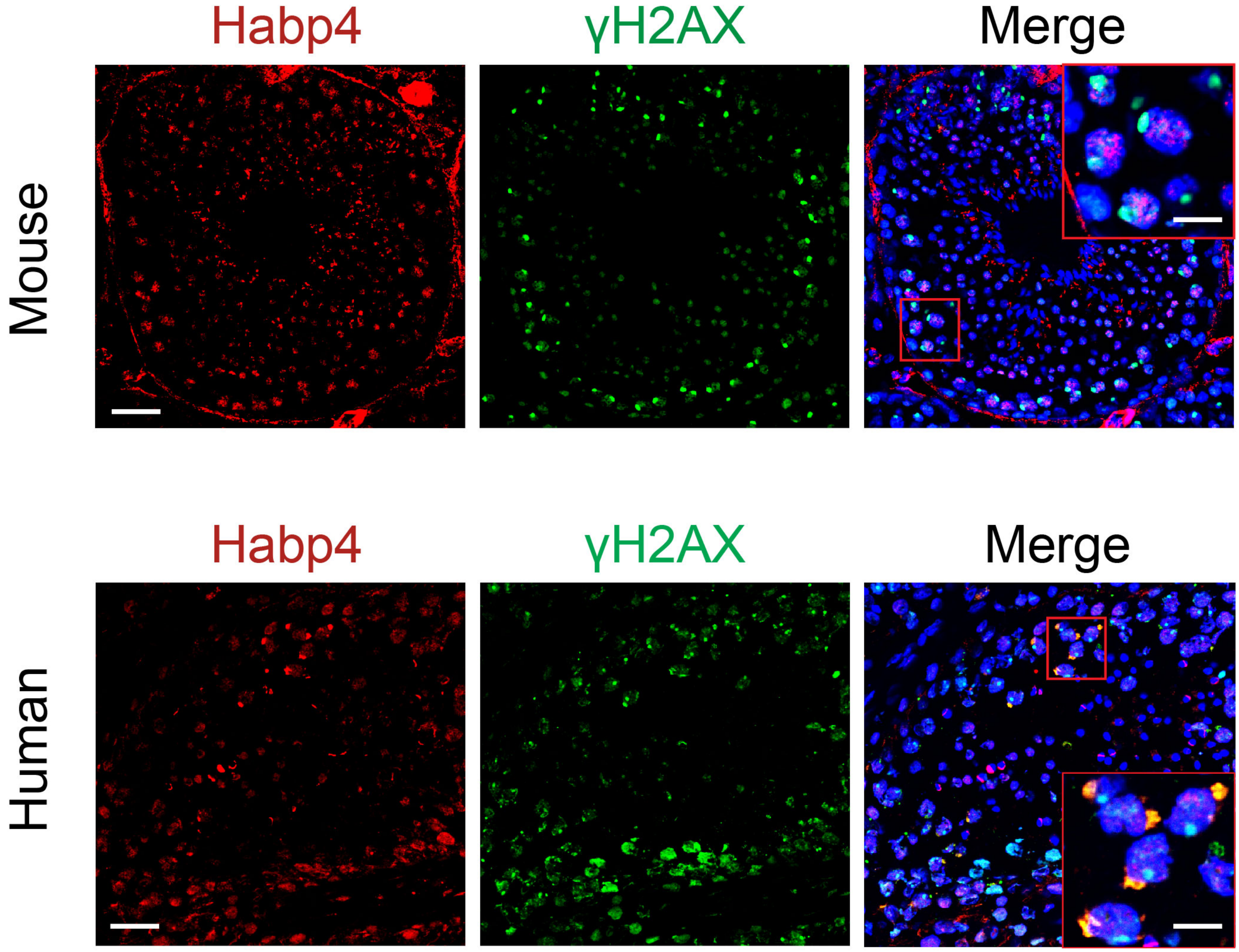


**Figure S4 Spatial localization of the Habp4 protein and γH2AX in mouse and human seminiferous tubules**
Habp4 co-localizes with the γH2AX, a molecular marker for monitoring DNA damage, in a subset of human spermatocytes, but not in the mouse. Scale bar for the mouse images, 40μm; 10 μm for the inset. Scale bar for the human images, 40μm; 15 μm for the inset.

### Figure S5

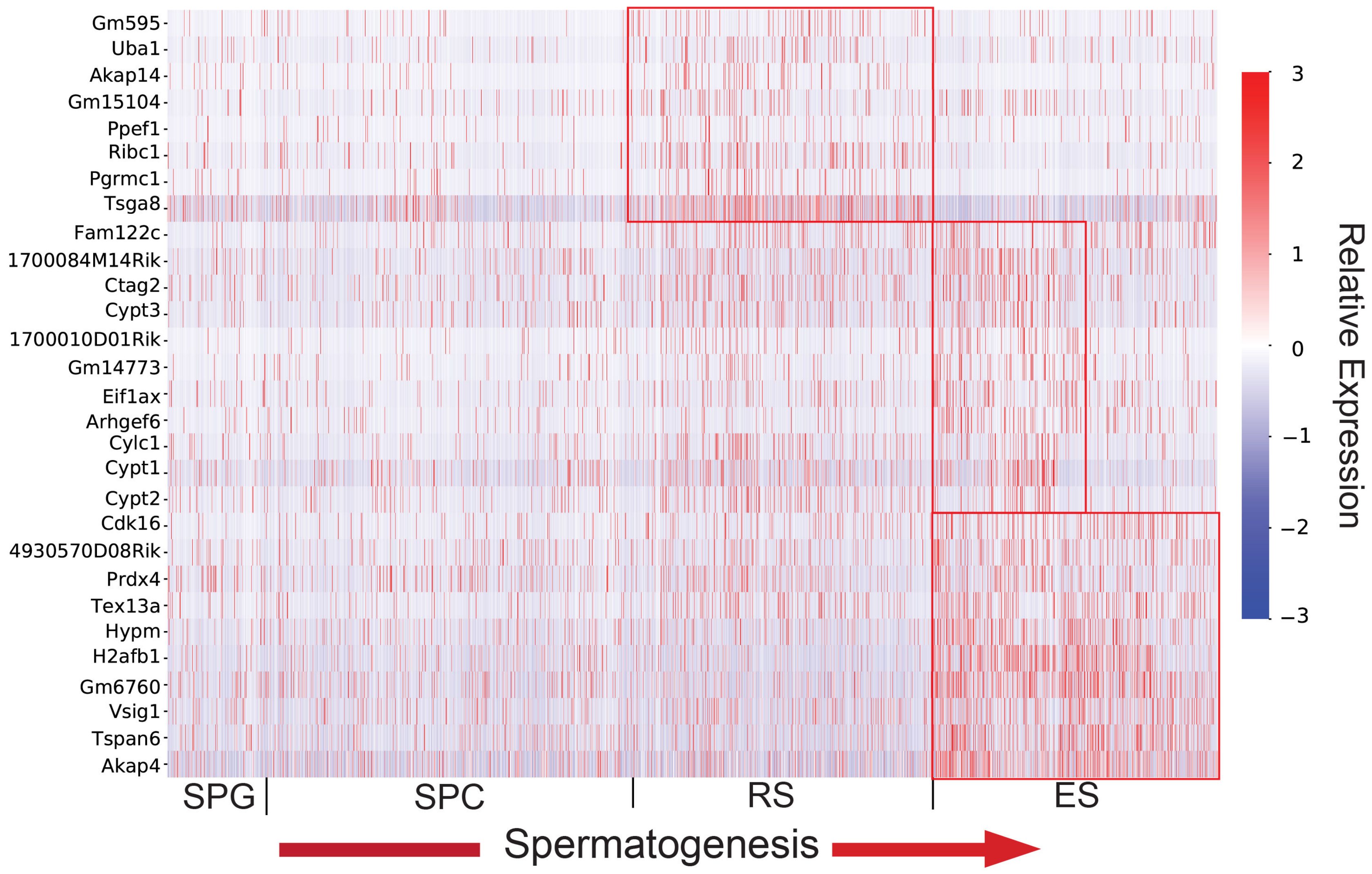


**Figure S5 Temporal dynamics of the meiotic sex chromosome inactivation (MSCI) escape genes during spermatogenesis**
A group of X chromosome MSCI escape genes exhibit three distinct temporal expression patterns (boxed) following meiosis.

### Figure S6

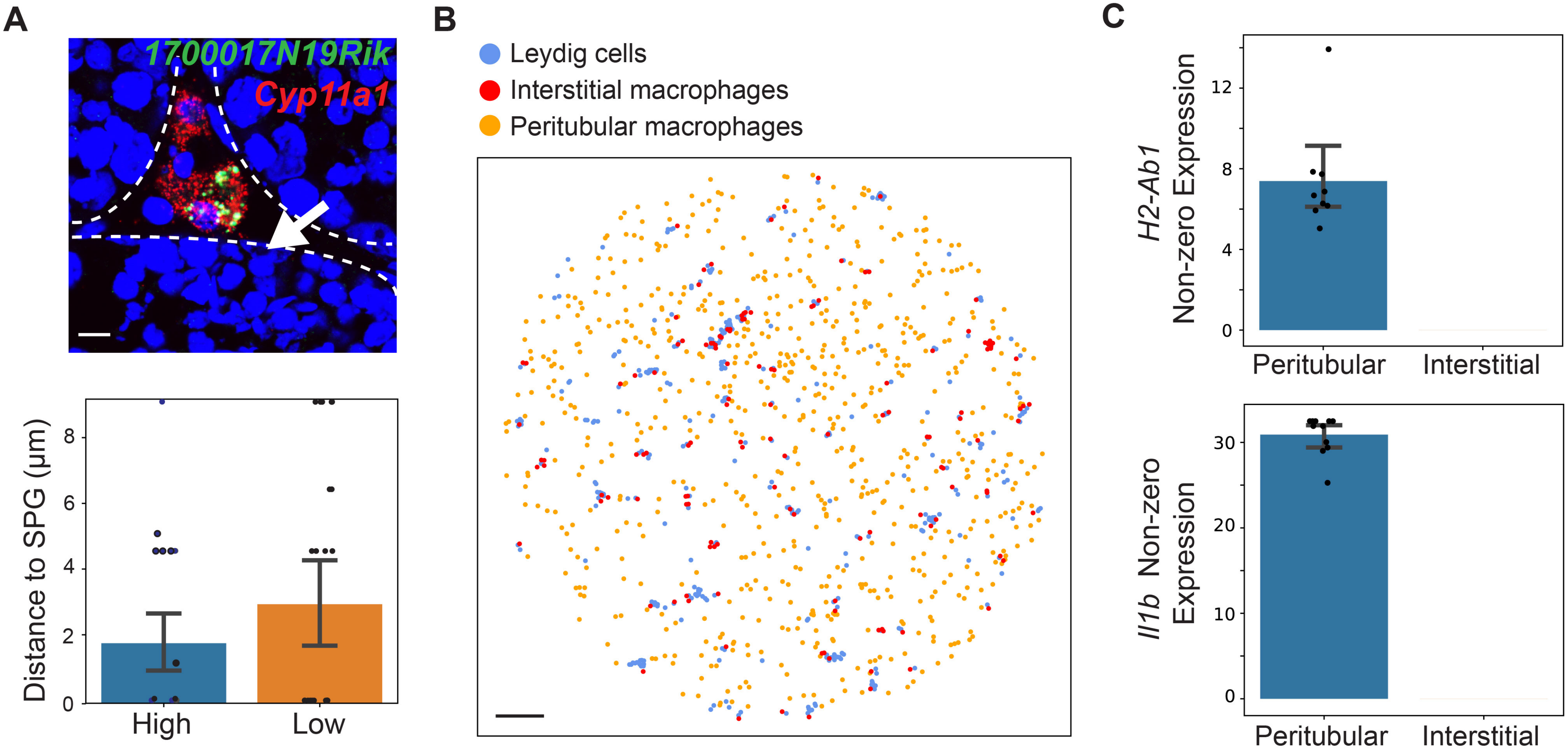


**Figure S6 Genes with stage-dependent expression patterns in Leydig cells and macrophages.**

(A) Upper panel: a representative image of a *1700017N19Rik*-expressing Leydig cells close to a spermatogonium (white arrow head). Scale bar, 10 µm. Lower panel: the comparison of the Euclidean distance between Leydig cells with high *1700017N19Rik* expression and the nearest spermatogonium and Leydig cells with low or no *1700017N19Rik* expression and the nearest spermatogonium. Two sided student t test, $p<0.05$.

(B) Spatial localizations of Leydig cells and macrophages in the spatial transcriptome atlas. The interstitial space is demarcated by the Leydig cell localizations. Scale bar, 300 µm.

(C) Genes *H2-Ab1* and *Il1b* are enriched in peritubular macrophages, but not in interstitial ones.

### Figure S7

**A**

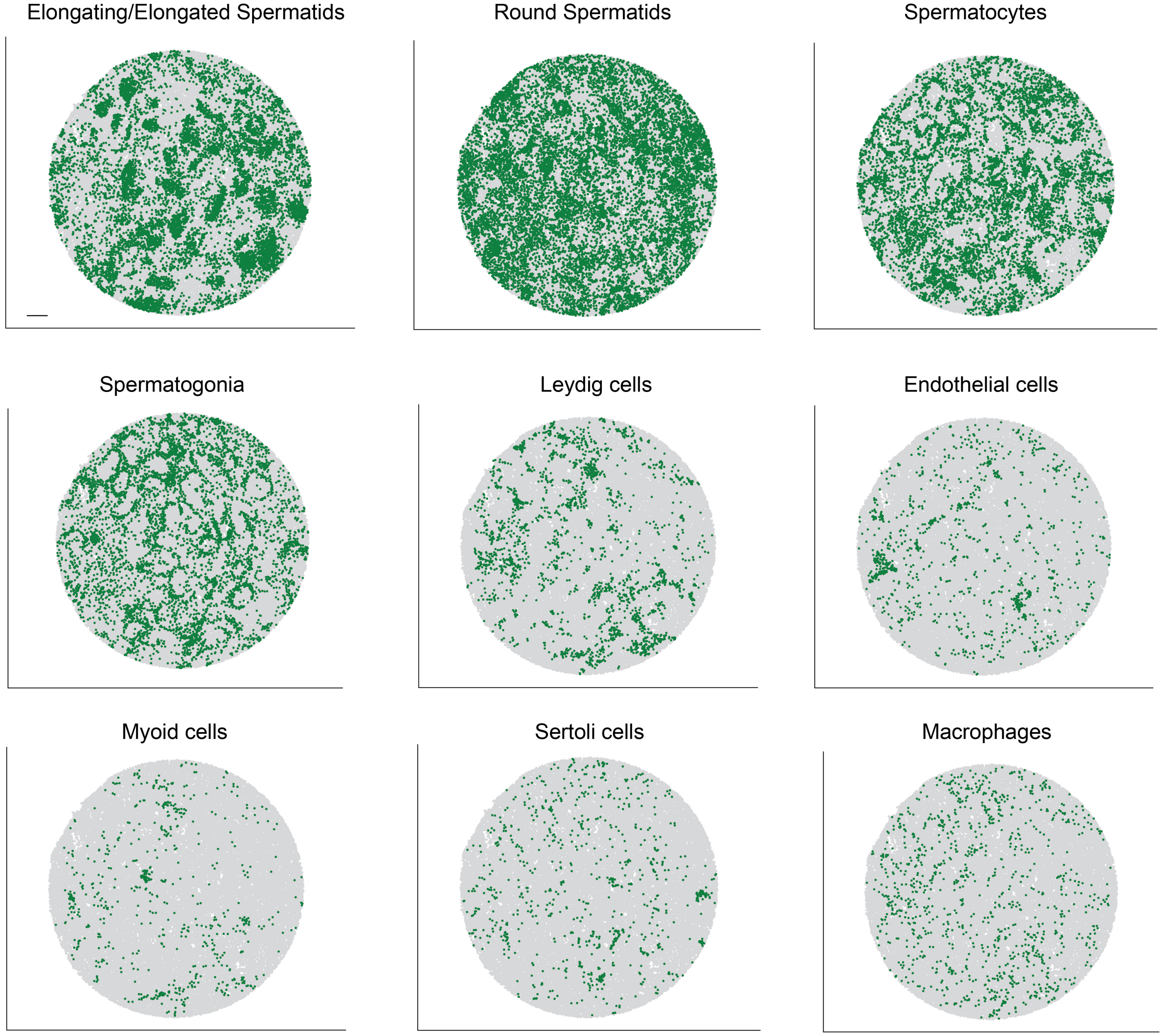


**B**

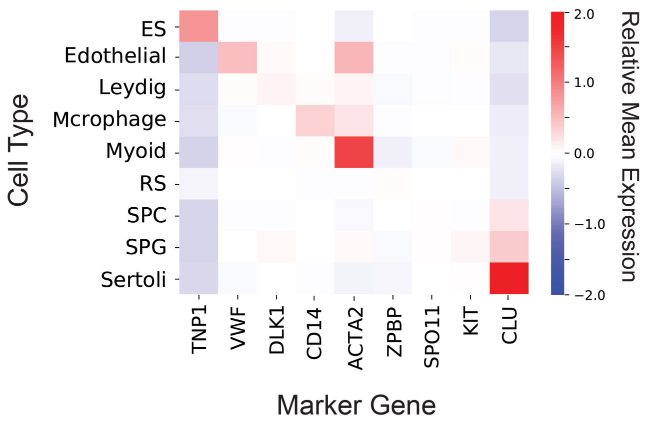


**Figure S7 Spatial mapping of individual human testicular cell types**

(A) The spatial localization of each testicular cell type. Scale bar, 300 μm.

(B) The mean gene expression matrix for cell type marker genes. ES, elongating/elongated spermatid; RS, round spermatid; SPC, spermatocyte; SPG, spermatogonium.

### Figure S8

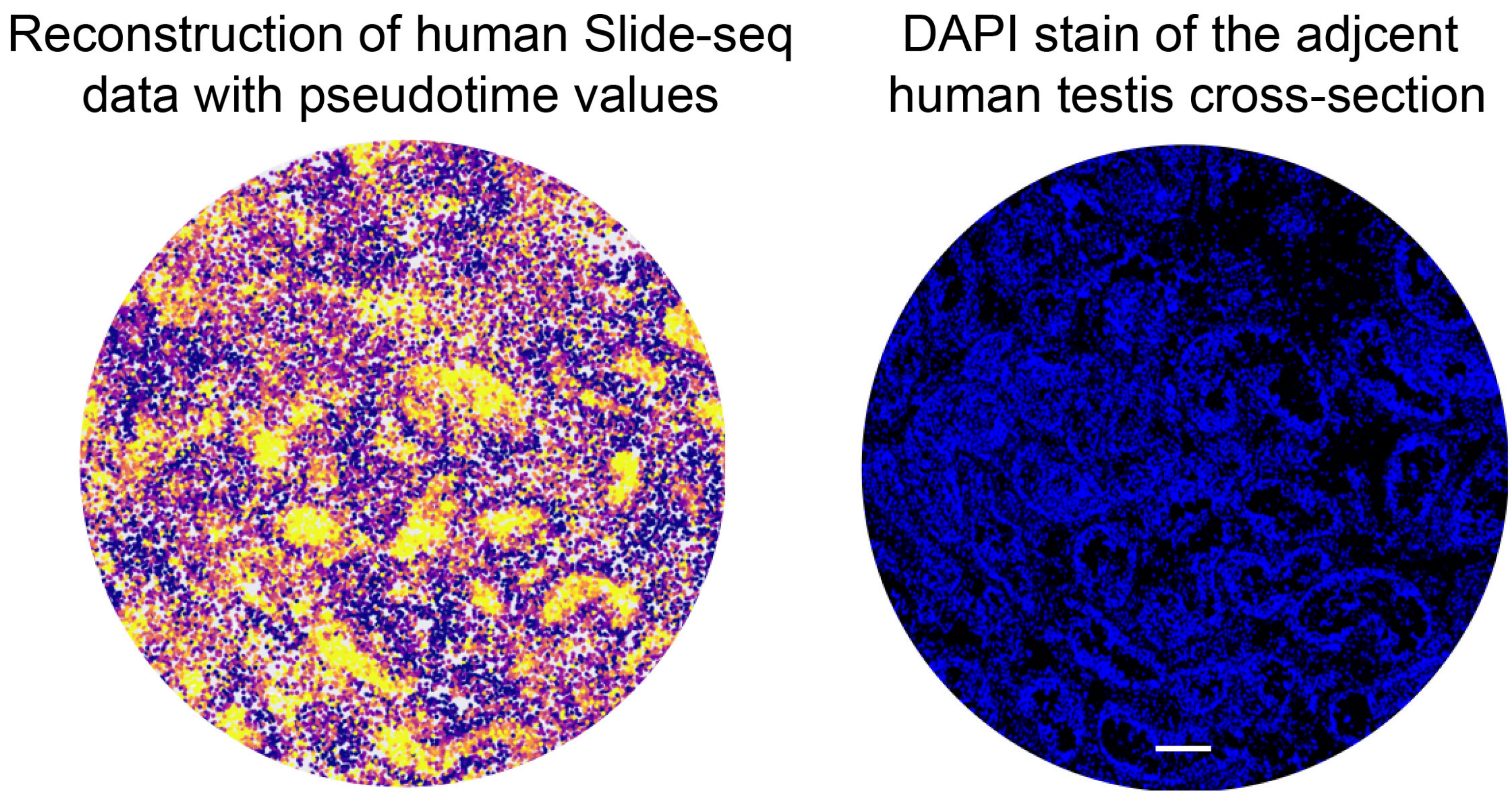


**Figure S8 The human testicular spatial atlas retains the morphological details of the profiled testis cross-section**
The pseudotime-reconstructed image of the human Slide-seq data shows consistent morphological features with the DAPI image of the adjacent testis cross-section. Scale bar, 200 μm.

### Figure S9

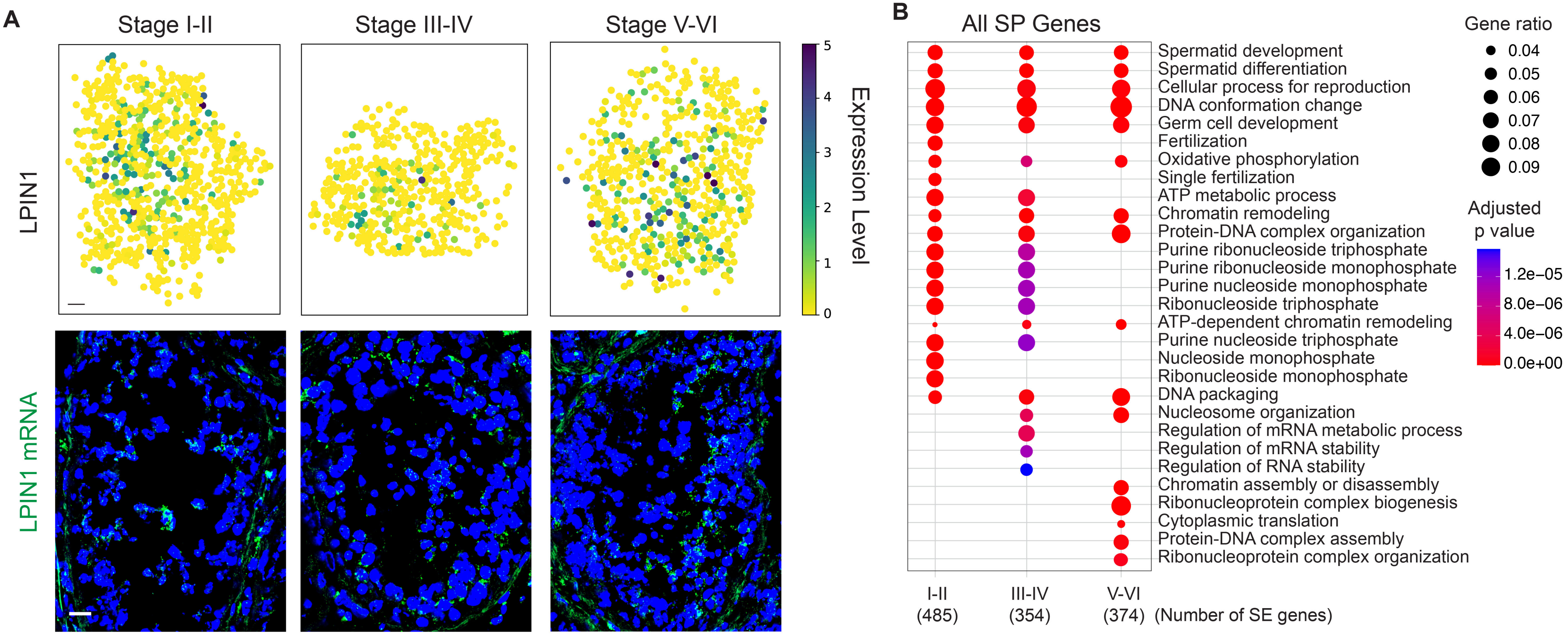


**Figure S9 Spatially patterned (SP) genes in the human seminiferous tubules**

(A) The spatial expression pattern of *LPIN1* revealed by both the spatial transcriptome atlas and smFISH. Scale bar, 30 μm for the digitally reconstructed seminiferous tubule images and 20 μm for the smFISH images.

(B) Gene ontology enrichment analysis on human SP genes from the three stage clusters.

### Figure S10

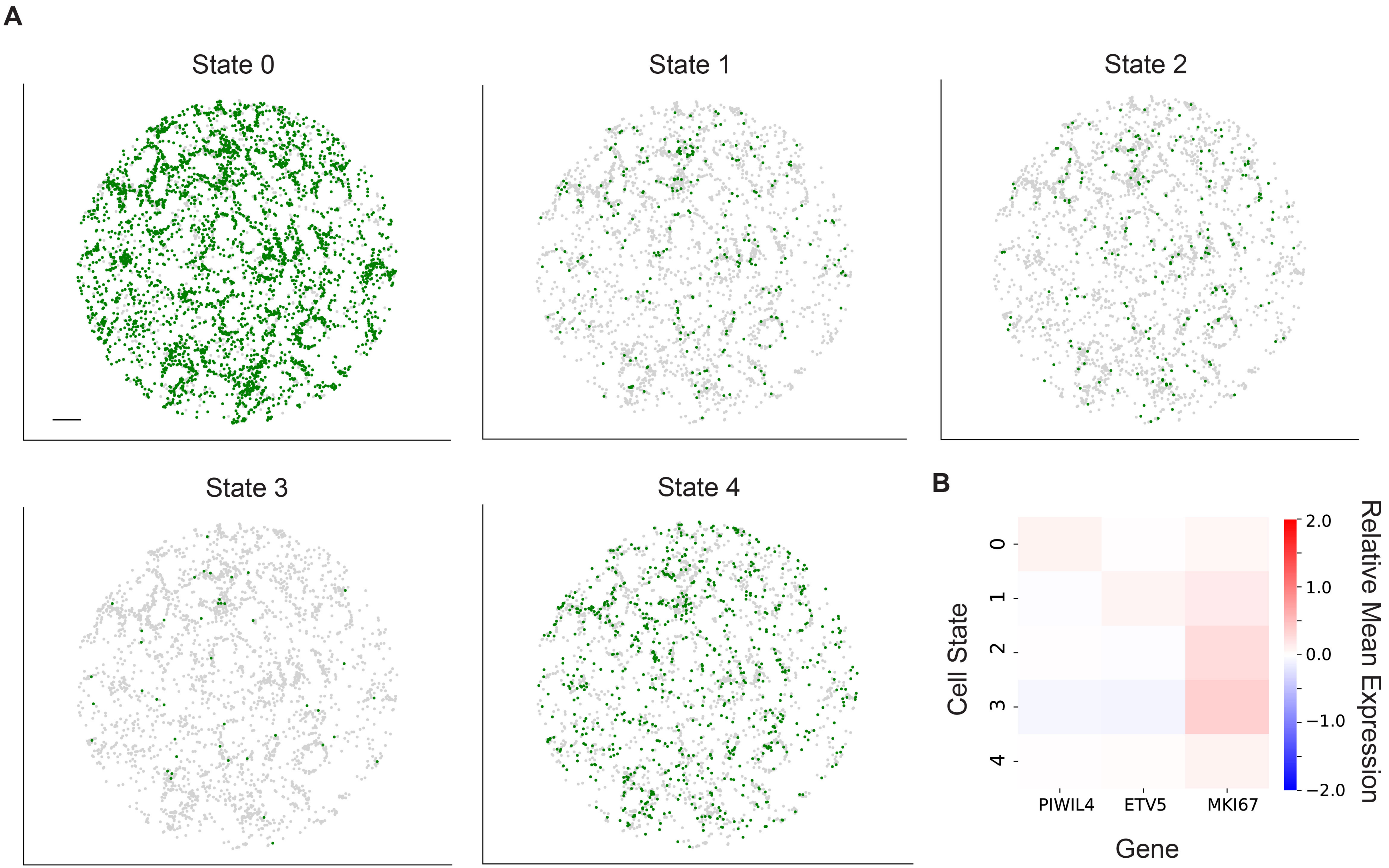


**Figure S10 Spatial mapping of the transcriptional states in the human spermatogonial population**
(A) The spatial localization of the five transcriptional states of human spermatogonia. Scale bar, 300 μm.
(B) The mean expressions of genes *PIWIL4*, *ETV5*, and *MKI167* in the five transcriptional states.

### Figure S11

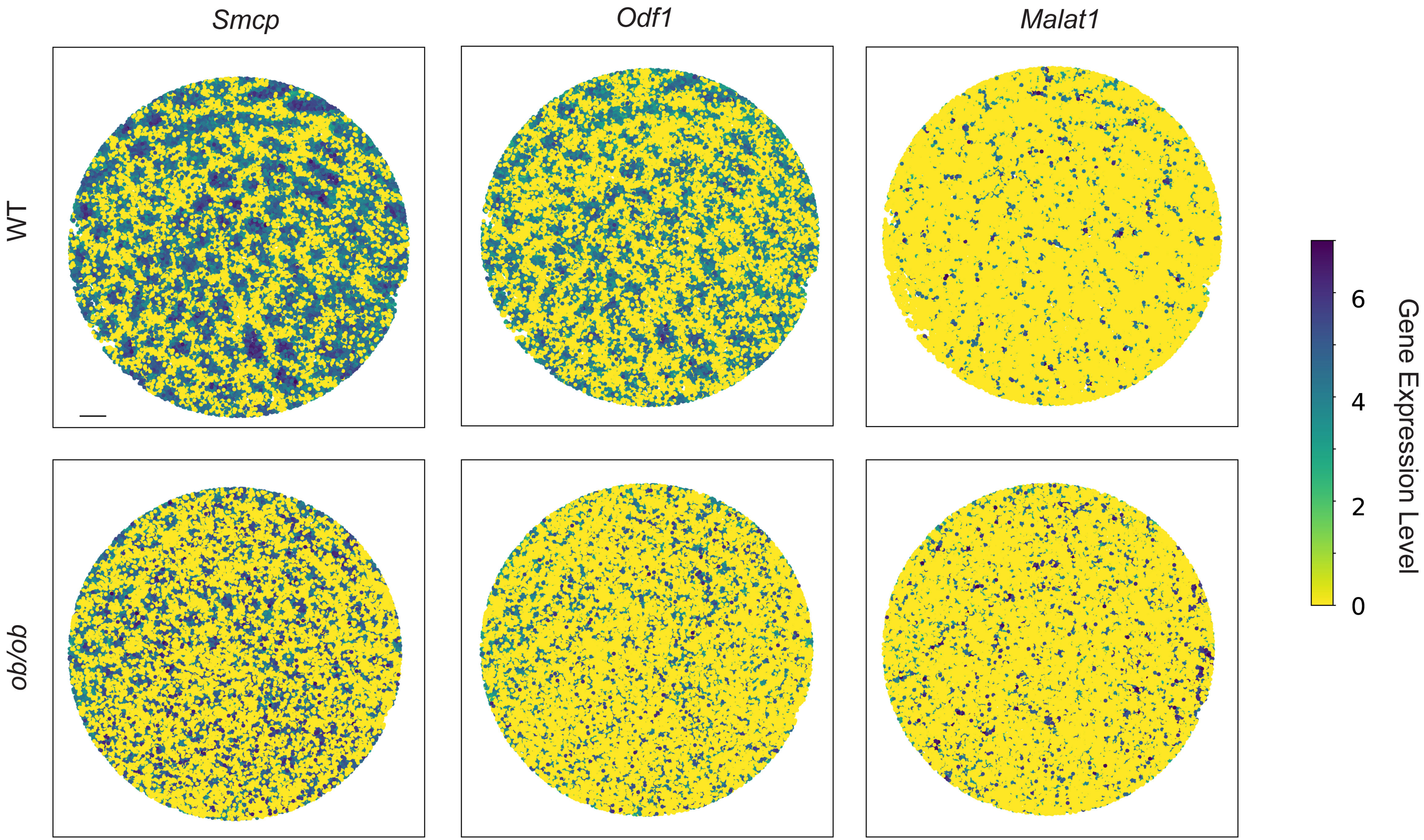


**Figure S11 Diabetes induces changes in spatial gene expression patterns**
The spatial expression pattern of genes *Smcp*, *Odf1*, *Malat1* in a representative WT and ob/ob seminiferous tubule, respectively. Scale bar, 300 μm.
